# Comparative genomics reveals factors associated with phenotypic expression of *Wolbachia*

**DOI:** 10.1101/2021.01.29.428792

**Authors:** Guilherme Costa Baião, Jessin Janice, Maria Galinou, Lisa Klasson

## Abstract

*Wolbachia* is a widespread, vertically transmitted bacterial endosymbiont known for manipulating arthropod reproduction. Its most common form of reproductive manipulation is Cytoplasmic incompatibility (CI), observed when a modification in the male sperm leads to embryonic lethality unless a compatible rescue factor is present in the female egg. CI attracts scientific attention due to its implications for host speciation and in the use of *Wolbachia* for controlling vector-borne diseases. However, our understanding of CI is complicated by the complexity of the phenotype, whose expression depends on both symbiont and host factors. In the present study, we perform a comparative analysis of nine complete *Wolbachia* genomes with known CI properties in the same genetic host background, *Drosophila simulans* STC. We describe genetic differences between closely related strains and uncover evidence that phages and other mobile elements contribute to rapid evolution of both genomes and phenotypes of *Wolbachia*. Additionally, we identify both known and novel genes associated with the modification and rescue functions of CI. We combine our observations with published phenotypic information and discuss how variability in *cif* genes, novel CI-associated genes and *Wolbachia* titer might contribute to poorly understood aspects of CI such as strength and bidirectional incompatibility. We speculate that high titer CI strains could be better at invading new hosts already infected with a CI *Wolbachia*, due to a higher rescue potential, and suggest that titer might thus be a relevant parameter to consider for future strategies using CI *Wolbachia* in biological control.

## INTRODUCTION

Endosymbiotic bacteria are associated with most insects and contribute to the biology and evolution of their hosts in multiple ways. Among the most well studied of these endosymbionts is *Wolbachia*, which infects 40% of all arthropods and several filarial nematodes (Zug and Hammerstein 2012). Although only one species is currently recognized, *Wolbachia* strains show considerable diversity and are organized in several supergroups (Lefoulon, et al. 2020; Lo, et al. 2007). *Wolbachia* can affect their hosts in different ways and are for example known to increase fecundity, longevity, fertility and provide protection against viruses (Fast, et al. 2011; Hedges, et al. 2008; Martinez, et al. 2015; Teixeira, et al. 2008). However, it is as a reproductive parasite that *Wolbachia* is best known, and its evolutionary success is often attributed to its efficacy in manipulating the host reproductive system to increase its own spread.

Since *Wolbachia* is vertically transmitted from mother to offspring via the egg, males represent a dead end for its transmission. Thus, three of the four known *Wolbachia* phenotypes that affect reproduction, feminization, parthenogenesis induction and male-killing, lead to more females than males being produced in infected host populations. However, the most common and well-studied phenotype, cytoplasmic incompatibility (CI), does not bias sex ratios. Instead, it is a form of sterility that results in embryonic mortality when an infected male mates with an uninfected female (unidirectional CI) or when a female and male carrying different and incompatible *Wolbachia* strains mate (bidirectional CI) (Werren, et al. 2008). As such, CI results in a reproductive advantage for infected females over uninfected females which leads to an effective spread of the symbiont in host populations. CI has so far been found in *Wolbachia* supergroups A and B, and the interest in the phenotype relates to both applied and basic research. Due to the ability to drive rapid symbiont spread and cause conditional sterility, CI is being used to introduce *Wolbachia* into mosquito populations in an attempt to reduce their numbers and to prevent the spread of dengue, Zika and chikungunya viruses (Flores and O’Neill 2018; Zheng, et al. 2019). From an evolutionary perspective, bidirectional CI is implicated in host speciation, as it creates a reproductive barrier between individuals that are infected with incompatible strains (Bordenstein, et al. 2001).

The phenotypic expression of CI is often described in terms of modification (*mod*) and rescue (*resc*) (Werren 1997). Modification occurs in the sperm of infected males before *Wolbachia* is shed. For offspring to be produced, the modified sperm have to fuse with a *Wolbachia*-infected egg containing a rescue factor. If the egg does not contain the correct rescue factor, development will halt when the embryo enters the first mitotic division as a result of asynchrony between the paternal and maternal chromosomes (Tram and Sullivan 2002). The *mod* and *resc* functions are independent, since some strains can rescue but not modify, and *Wolbachia* strains can be classified based on their ability to exert them. Although all variations exist, most strains seem to either modify sperm and rescue their own modification (*mod^+^ resc^+^*) or neither modify nor rescue (*mod^−^ resc^−^*) (Poinsot, et al. 2003; Zabalou, et al. 2008).

Recently, the phage-associated genes *cifA* and *cifB* were shown to play a major role in the CI phenotype, although their exact functions in terms of *mod* and *resc* are still debated. While *cifB* is undoubtedly linked to *mod*, it is not clear if *cifA* is involved only in *resc* or both *mod* and *resc* (Beckmann, et al. 2019; Shropshire and Bordenstein 2019). In the latter hypothesis, known as the two-by-one genetic model of CI, both *cifA* and *cifB* are required for causing modification, while *cifA* alone performs rescue when expressed at an appropriate level (Shropshire and Bordenstein 2019; Shropshire, et al. 2018). Homologs of *cifA* and *cifB* have been identified in various *Wolbachia* strains as well as in a few other Rickettsiaceae, and they are classified into five Types (I-V) according to phylogenetic analyses (Lindsey, et al. 2018; Martinez, et al. 2020). The different *Wolbachia cif* types show considerable variation in sequence length and predicted protein domains, but are all expected to perform CI-associated functions (LePage, et al. 2017; Martinez, et al. 2020). Accordingly, a strong correlation exists between strains carrying *cif* genes that are predicted to be functional and those known to induce and rescue CI (LePage, et al. 2017; Martinez, et al. 2020). There is also a general trend for strains carrying phylogenetically related *cif* genes to be compatible with each other, although some exceptions have been observed (Shropshire, et al. 2020b). Experimental evidence for the ability of *cifA*-*B* to cause and rescue CI exists for Type I genes from *Wolbachia* strain *w*Mel of *D. melanogaster* and Types I and IV from *Wolbachia* strain *w*Pip of *C. pipiens*. (Chen, et al. 2019; LePage, et al. 2017). Interestingly, the *mod* function of Type I is associated with a deubiquitylase domain while that of Type IV is linked to a nuclease domain (Chen, et al. 2019; LePage, et al. 2017). This suggests that distinct molecular mechanisms of CI might exist (Lindsey, et al. 2018; Martinez, et al. 2020), and recent evidence points to multiple domains likely being involved in both *mod* and *resc* functions (Shropshire, et al. 2020a). Furthermore, the *cif* genes alone do not explain all *mod* and *resc* phenotypic variations observed in different systems, especially with regard to CI strength and bidirectional incompatibility between strains (Shropshire, et al. 2020b). This is observed, for example, in the *w*Mel strain which only has Type I *cifA-B* but can partially rescue the modification of *w*Ri, which only has functional *cifB* of Type II (Charlat, et al. 2004; Zabalou, et al. 2008). Such cases suggest that CI phenotypic expression is also modulated by other genes and factors such as *Wolbachia* titer and host localization (Shropshire, et al. 2020b).

Several mechanistic models have been proposed to explain CI *mod* and *resc* (Beckmann, et al. 2019; Bossan, et al. 2011; Poinsot, et al. 2003; Shropshire, et al. 2019). Poinsot, et al. (2003) evaluated three different models and concluded that the “lock-and-key” best fit the knowledge at the time. This model suggests that the *mod* factor puts a lock on the paternal chromosome and a matching key, the *resc* factor, has to be present in the egg in order for the paternal chromosome to enter mitosis. The model requires that the *mod* and *resc* functions are unique and encoded by separate bacterial genes. Later, Bossan, et al. (2011) combined the qualitative lock-and-key model with added quantitative parameters such as timing and expression, making the model fit better with observations regarding the ability of several *Wolbachia* strains to induce CI and rescue each other’s modification. Currently, the Toxin-Antidote (TA) and Host-Modification (HM) models are the main mechanistic hypotheses for CI (Beckmann, et al. 2019; Shropshire, et al. 2019). The TA model is generally similar to the lock and key and suggests that *Wolbachia* releases a toxin in the male sperm which must be counteracted by an appropriate antidote in the female egg (Beckmann, et al. 2019; Hurst 1991). The HM model, on the other hand, postulates that *Wolbachia* alters a host product in the male sperm which leads to embryonic mortality unless the modification is reversed by a *Wolbachia* factor in the egg (Shropshire, et al. 2019).

Independently of its mechanistic model, it is clear that the phenotypic expression of CI as well as of other *Wolbachia* phenotypes depends not only on symbiont factors but also on the host’s genetic background. The same *Wolbachia* strain can, for example, cause male-killing in one host and CI in another (Jaenike 2007; Sasaki, et al. 2005). One strain can also induce different strengths of CI when transferred between host species. This is seen when the *w*Mel strain that naturally infects *Drosophila melanogaster* and the *w*Ri strain that naturally infects the related species *D. simulans,* are transferred to each other’s natural host. In its natural host, *w*Mel induces up to 30% embryonic mortality, whereas in *D. simulans* it causes almost 100% CI (Poinsot, et al. 1998). The opposite effect can be seen for *w*Ri, that causes almost 100% embryonic mortality in its natural host but only around 30% in *D. melanogaster* (Boyle, et al. 1993). Similarly, the two strains *w*Tei and *w*MelPop induce no or weak CI in their natural hosts (*D. teissieri* and *D. melanogaster,* respectively), but almost 100% embryonic mortality when transferred into *D. simulans* (McGraw, et al. 2001; Zabalou, et al. 2008). These examples also show that *D. simulans* is a permissive host in which many *Wolbachia* strains induce stronger CI in comparison to other *Drosophila* species. The permissiveness of *D. simulans* is also reflected in the variety of *Wolbachia* strains that naturally infect this species, at least five, and in the many successful experimental transfers of *Wolbachia* from other hosts into *D. simulans* (Merçot and Charlat 2004). As a result, *D. simulans* became an important model for CI studies and phenotypic comparisons between *Wolbachia* strains (Martinez, et al. 2015; Merçot and Charlat 2004; Zabalou, et al. 2008).

In this paper, we investigate *Wolbachia* genome evolution with a focus on CI-associated genes by using five newly sequenced (*w*San, *w*Yak, *w*Tei, *w*Au and *w*Ma) and four previously available (*w*Ri, *w*No, *w*Ha and *w*Mel) complete *Wolbachia* genomes. All nine strains have known *mod* and *resc* phenotypes in the *D. simulans* STC host background (Zabalou, et al. 2008), and five of them naturally infect *D. simulans.* Among these five, three are *mod^+^resc^+^* (*w*Ri, *w*Ha and *w*No) and show variable CI strength, while two (*w*Ma and *w*Au) do not induce CI (*mod^-^*). The non-CI inducers still differ in their rescue properties, with *w*Au incapable of rescue (*resc^-^*) while *w*Ma can rescue the modification of *w*No (*resc^+^*). Three other strains, *w*San, *w*Yak and *w*Tei (hereafter referred to as *w*SYT when mentioned collectively), naturally infect the species of the *Drosophila yakuba* group, *D. santomea*, *D. yakuba*, and *D. teissieri*, respectively. These are closely related and cause no to low CI in their natural hosts but show different phenotypes after being transferred to *D. simulans.* In the new host*, w*San and *w*Yak continue causing no or low CI while *w*Tei induces a strong incompatibility (Cooper, et al. 2017; Martinez, et al. 2015; Zabalou, et al. 2008; Zabalou, et al. 2004). The *w*SYT strains also differ in their infection titer and in their compatibility with other strains, as *w*Tei has significantly higher titer than *w*SY (Martinez, et al. 2015) and is capable of rescuing the modification of *w*Mel while *w*SY are not (Zabalou, et al. 2008).

Thus, our dataset focuses on a single host and includes closely related *Wolbachia* strains with distinct phenotypes, the *w*SYT and *w*No-*w*Ma. This creates a unique opportunity for identifying *Wolbachia* factors associated with specific traits of each strain, among which *mod*, *resc* and CI strength. Most of our strains also have previously described non-reproductive phenotypes in *D. simulans* STC, which allows us to analyze factors associated with cell and tissue tropism (Toomey and Frydman 2014; Toomey, et al. 2013; Veneti, et al. 2004), protection against viruses (Martinez, et al. 2014), fecundity and female lifespan (Martinez, et al. 2015). In particular, we focus on the very closely related genomes of *w*SYT and investigate their most variable regions in order to identify genetic elements correlated with the phenotypic differences between them. Additionally, we investigated mutations that have occurred after transfection, and that might be due to changes in host selection, by sequencing multiple independently transfected line of the *w*SYT strains from *D. simulans* STC as well as a few from their native hosts.

Our results identify unique genetic features of our strains and highlight the importance of mobile elements to *Wolbachia* evolution, uncovering lateral gene transfers between *Wolbachia* strains as well as between *Wolbachia* and other organisms. By screening for CI-associated genes we recover the *cif* genes and identify novel candidate genes potentially associated with *mod* and *resc.* We compare our results with published phenotypic information for our strains and discuss how CI-associated genes as well as symbiont titer may influence CI strength and compatibility between *Wolbachia* strains. Overall, this study contributes to further understanding of the CI phenotype and the genomic flexibility that allows *Wolbachia* to accumulate genetic changes with potentially major effects on both host and symbiont within short time scales.

## RESULTS

### Genome features and strain relationship

The five *Wolbachia* genomes that were completely sequenced in this study, *w*San, *w*Yak, *w*Tei, *w*Au and *w*Ma, are circular and range in size between 1.27 and 1.41 Mbp (Table 1). Their genome size and features are similar to the four previously sequenced *Wolbachia* genomes used in our comparative analyses (Ellegaard, et al. 2013; Klasson, et al. 2009b; Sutton, et al. 2014; Wu, et al. 2004) as well as to many other sequenced *Wolbachia* genomes (Klasson, et al. 2008; Newton, et al. 2016; Sinha, et al. 2019).

**Table 1.** Genome features of *Wolbachia* genomes.

| Genomes | wMel | wAu | wSan | wYak | wTei | wRi | wHa | wMa | wNo |
| --- | --- | --- | --- | --- | --- | --- | --- | --- | --- |
| Supergroup | A | A | A | A | A | A | A | B | B |
| Genome size (Mbp) | 1.27 | 1.27 | 1.41 | 1.39 | 1.35 | 1.45 | 1.30 | 1.27 | 1.30 |
| GC (%) | 35.2 | 35.2 | 35.2 | 35.2 | 35.2 | 35.2 | 35.1 | 34.0 | 34.0 |
| Genes | 1199 (1011) | 996 | 1120 | 1102 | 1069 | 1150 | 1009 | 1006 | 1042 |
| Coding density | 0.80 (0.76) | 0.76 | 0.77 | 0.76 | 0.77 | 0.80 | 0.78 | 0.80 | 0.81 |
| Avg. gene length (bp) | 851 (958) | 963 | 963 | 965 | 968 | 976 | 1001 | 1015 | 1012 |
| Pseudogenes | 74 (111) | 122 | 118 | 115 | 116 | 114 | 96 | 89 | 91 |
| Phage WO (%) | 8 | 9 | 14 | 13 | 10 | 8 | 8 | 6 | 8 |
| ANK<sup>§</sup> (%) | 3 | 3 | 3 | 3 | 2 | 3 | 3 | 6 | 7 |
| Pseudogenes (%) | 6 | 10 | 9 | 9 | 11 | 8 | 7 | 7 | 7 |
| Repeats<sup>*</sup> (%) | 9 | 10 | 17 | 15 | 11 | 23 | 10 | 4 | 4 |

<sup>§</sup> Ankyrin repeat domain proteins <sup>\*</sup> Repeats longer than 300 bp with higher than 95% identity ( ) Numbers in parenthesis were obtained after manual curation of the wMel original annotation

The *w*Au genome sequenced here differs by five SNPs and five indels from the *w*Au genome published by Sutton, et al. (2014). All five SNPs are present in intergenic regions, of which four are in repeats. The five indels are all present in repeat regions, two of which cause pseudogenization of mobile elements in the genome published by Sutton, et al. (2014). Even though the sequences of the two *w*Au genomes themselves are very similar, the annotation differs considerably, as we have used a different annotation pipeline followed by manual curation. For consistency, all of our analyses were done using the *w*Au genome sequence and annotation presented in this study.

In order to further increase the consistency of the annotations between our compared genomes, we also manually curated the *w*Mel annotation (numbers present in parenthesis in Table 1). Although *w*Mel is closely related to *w*SYT and *w*Au, the coding density in the original annotation of wMel is higher and the average gene length is shorter than in the *w*SYT and *w*Au genomes (Table 1). These differences are mostly due to dissimilarities in pseudogene annotation (6% vs. ~10%) and were alleviated by our manual curation. Furthermore, since one of our goals with the study is to identify candidates associated with phenotypic differences in a controlled host background, we also changed the gene sequences of *w*Mel in accordance with a *w*Mel strain that we sequenced after transinfection to *D. simulans* (see results below).

To establish a robust phylogeny between the nine *Wolbachia* strains (Table 1), we clustered their proteomes and used the resulting 714 single copy orthologous genes for phylogenetic reconstruction. A maximum likelihood tree based on the concatenated alignment of these genes showed 100% bootstrap support for all nodes (Figure 1). Although the branch lengths are very short, it is clear that *w*San and *w*Yak are most closely related followed by *w*Tei and that the *w*SYT genomes group together with *w*Mel to the exclusion of *w*Au. This result is in agreement with Cooper, et al. (2019) and in contrast to Zabalou, et al. (2008), who found *w*Au branching closest to *w*SYT.

**Figure 1.**
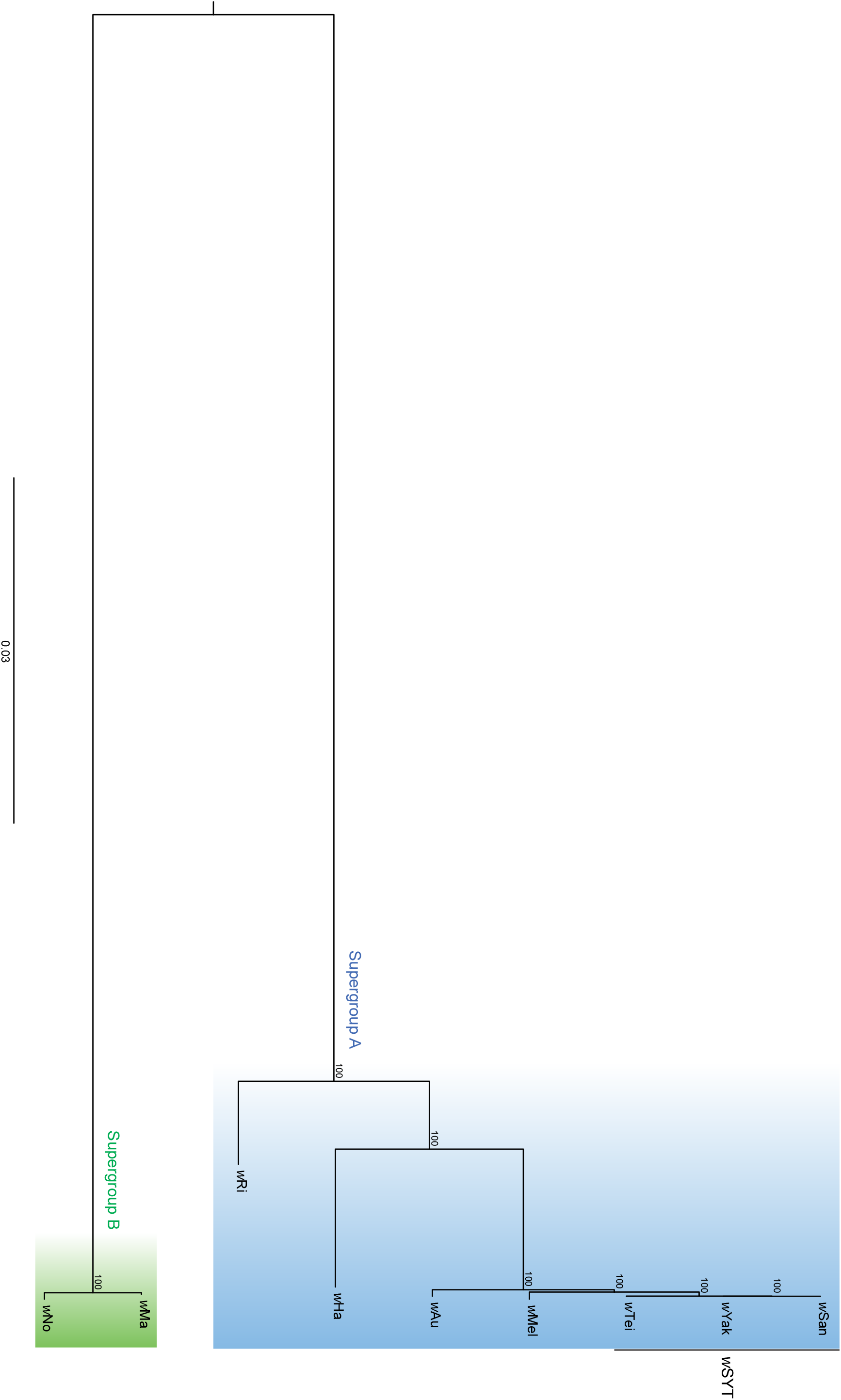
Phylogenetic relationships of the nine *Wolbachia* genomes included in this study. Maximum likelihood tree based on the concatenated alignment of 714 orthologous single copy genes. All nodes have 100% bootstrap support, as indicated by an asterisk.

### Mutations after transfer to *D. simulans*

In order to investigate what mutations might have occurred after transfer to a new host, we sequenced DNA from multiple independent *Drosophila* lines for our supergroup A *Wolbachia* strains (Table 1). Three separate *Drosophila* lines infected with *w*Tei and *w*Yak and two lines infected with *w*Au and *w*San (Table S1) were sequenced, the reads were mapped against their respective closed genome and variants were called. Additionally, one *D. simulans* line transinfected with *w*Mel was sequenced and compared to the publicly available *w*Mel genome (Wu, et al. 2004). As a control, we also run the same pipeline with the reads from the exact same strain used for producing the reference.

In *w*Au, *w*Tei and *w*Yak, there were SNPs called between the reference and the Illumina reads used to create it (Table S2). In all those positions, we found discrepancies between the PacBio and Illumina reads and we chose to call the sequence according to the PacBio reads. However, all such SNPs are present in intergenic regions, so their impact on our analyses is minimal.

In the comparisons between *Wolbachia* genomes from the same strain but different *Drosophila* lines, we found a few SNPs located mostly in intergenic regions (Table S2). Only in two of the comparisons did we observe mutations that would likely alter the function of a protein.

First, the *w*Yak strain sequenced from its natural host *D. yakuba* had an indel that causes a frameshift in a gene that codes for a permease. Since it is a loss-of-function mutation that is not present in the transinfected line nor in the published draft assembly of *w*Yak from *D. yakuba* (GCA_005862115.1), it seems most likely that this mutation occurred in our sequenced line from *D. yakuba* and not when transferring *w*Yak to *D. simulans*.

Second, in the *w*Mel strain sequenced from *D. simulans*, we found indels in four genes, all coding for hypothetical proteins. We believe that all four might represent errors or possibly mutations that occurred in the published *w*Mel genome (Wu, et al. 2004) rather than after transfer to *D. simulans*. Three of the indels restores the frame so that two short ORFs become one long (WD1043-WD1044, WD1215-WD1216, WD1231-WD1232), possibly leading to functional restoration of the affected proteins. The last indel puts WD1155-WD1156 in the same frame, creating a new long putative gene that contains an in-frame stop codon. The resulting sequence is similar to other A-supergroup genomes sequenced in this study that also contain the same in-frame stop codon (*w*SYT and *w*Au) and hence we believe that this might also be an error in the *w*Mel genome or mutation in the sequenced *w*Mel strain.

Overall, we didn’t identify any parallel mutations, i.e. mutations occurring in the same gene, between the genomes that have been transinfected into *D. simulans*, indicating that there isn’t a strong selection pressure on any particular protein as a result of transfer to the new host background.

### Genomic variation between close relatives

Among the nine genomes compared in this study, there are two clades of very closely related *Wolbachia* strains, *w*SYT plus *w*Mel and *w*Au (hereafter SYTMA), and *w*No and *w*Ma (hereafter NoMa). To estimate the overall level of divergence between the *Wolbachia* strains within each of these clades, Illumina reads from each strain within SYTMA and NoMa were mapped against each genome within the clade and variants were called. Variants for each pair of strains were calculated twice, since the numbers vary slightly depending on which of the two genomes was used as reference (Table 2, 3). Using the resulting SNP variants, we calculated the number of synonymous and non-synonymous mutations and compared them to the frequency of non-synonymous sites in the genomes, estimated to be 76% in the 714 single copy orthologs between all nine genomes. Based on their genomic location, we also classified the variants, both SNPs and indels, into genic, phage, and intergenic (Table S3). Additionally, we analyzed gene content differences between the most closely related genomes, *w*SYT and NoMa, and proteins that were uniquely present in the genomes of the SYTMA clade.

**Table 2.** Total number of SNPs between genomes of SYTMA and NoMa.

**A. SNPs between SYTMA genomes**
| Genomes<sup>§</sup> | wAu | wMel | wSan | wTei | wYak |
| --- | --- | --- | --- | --- | --- |
| wAu | - | 3199 | 2666 | 3202 | 2929 |
| wMel | 2914 | - | 2108 | 2636 | 2112 |
| wSan | 2924 | 2898 | - | 65 | 32 |
| wTei | 3039 | 2871 | 68 | - | 65 |
| wYak | 2742 | 2666 | 37 | 65 | - |

<sup>§</sup> Illumina reads from strains shown on each row were mapped to the genomes indicated on each column

**B. SNPs between NoMa genomes**
| Genomes<sup>§</sup> | wNo | wMa |
| --- | --- | --- |
| wNo | - | 1095 |
| wMa | 983 | - |

<sup>§</sup> Illumina reads from strains shown on each row were mapped to the genomes indicated on each column

**Table 3.** Total number of indels between genomes of either the A and B supergroups.

**A. Indels between SYTMA genomes**
| Genomes<sup>§</sup> | wAu | wMel | wSan | wTei | wYak |
| --- | --- | --- | --- | --- | --- |
| wAu | - | 401 | 383 | 457 | 428 |
| wMel | 348 | - | 234 | 308 | 236 |
| wSan | 354 | 317 | - | 7 | 4 |
| wTei | 357 | 276 | 6 | - | 12 |
| wYak | 316 | 260 | 4 | 12 | - |

<sup>§</sup> Illumina reads from strains shown on each row were mapped to the genomes indicated on each column

**B. Indels between NoMa genomes**
| Genomes<sup>§</sup> | wNo | wMa |
| --- | --- | --- |
| wNo | - | 87 |
| wMa | 90 | - |

<sup>§</sup> Illumina reads from strains on each row were mapped to the genomes indicated on each column

#### Variation between the wSYT strains

We found the three *w*SYT genomes to be extremely similar, differing only by 32-68 SNPs and 4-12 indels, thus making them 99,995% identical to each other in sequence (Table 2A, 3A). Using the pipeline described above, we observed that mutations were slightly underrepresented in the prophage WO regions, with only ca 5% of the total number of SNPs even though phage WO regions make up ca 10% of the genomes. Additionally, the genic SNP pattern indicated that purifying selection might not have had enough time to act on these mutations, as the frequency of non-synonymous mutations (75-80%) was close to neutrality (76%). Even so, there is an apparent overrepresentation of substitutions in intergenic regions (50-60%).

When analyzing our protein clusters, we didn’t find any cluster that was unique to either one of the three *w*SYT genomes, further emphasizing the close relationship between these three *Wolbachia* strains. Only five protein clusters were present in *w*SY to the exclusion of *w*Tei, even though the genomes of *w*SY are clearly larger than *w*Tei. Three of the five clusters contain phage WO proteins; three copies of a putative phage tail protein and the baseplate assembly proteins GpJ and GpW (Table S4). Only one protein is unique to *w*SY among our nine clustered proteomes and it is found as a pseudogene in *w*Tei (located in the *Island* described below). We also identified clusters with variation in copy number between *w*SYT genomes. A total of 31 clusters contain more copies in *w*SY than in *w*Tei, of which 29 are associated with phage WO and one is a putative non-WO phage terminase. The higher number of prophage proteins in the *w*SY genomes agrees with the larger proportion of phage detected in their genomes (Table 1) and also explains the larger genome sizes of *w*SY compared to *w*Tei. Only three protein clusters have more copies in *w*Tei than in *w*SY, a transposase of the IS4 family, a Group II intron associated reverse transcriptase, and the *cifB* gene wTei_05160, discussed in more detail in the next section.

Additional variation between the three *w*SYT genomes exists in the copy number of an IS-element of the IS5 family (23 copies in *w*Tei, 19 in *w*Yak and 18 in *w*San) as well as in the pseudogenization of two reverse transcriptase and the phage WO associated major tail sheath protein in *w*Yak.

Finally, in contrast to the very low number of mutations and few gene content differences, we observed that the gene order is highly variable between the *w*SYT genomes (Figure S1).

#### Variation between the NoMa strains

In the comparison between the NoMa genomes, we called approximately 1000 SNPs and 90 indels, making them 99,925% identical in sequence (Table 2B, 3B). Looking at the distribution of SNPs across the genomes, we found that the prophage WO SNPs were very slightly overrepresented (10%).

We identified 16 protein clusters present in *w*Ma to the exclusion of *w*No (12 are pseudogenes and 4 are completely absent). In *w*No, we identified 38 clusters that were absent from the *w*Ma genome (14 are pseudogenes and 24 are completely absent) (Table S4). These clusters are largely made up of a few categories of proteins, Ankyrin repeat (ANK) containing proteins (9), hypothetical proteins (21) and phage WO proteins (12). Very few of the genes that differ between the NoMa genomes are unique to either *w*Ma or *w*No. Instead, they are also present in other *Wolbachia* genomes (either in the ones included in our clustering or in the nr database). Only one gene in *w*Ma, encoding a hypothetical protein (wMa_00720) was found to be unique among currently sequenced *Wolbachia* genomes. In *w*No, two large ANK proteins (wNo_01030 and wNo_10640) are unique as they only have low similarity, low coverage hits to other *Wolbachia* genomes.

#### Variation in the SYTMA clade

The more distant genomes of the SYTMA clade have about 99,8% overall sequence identity (i.e. *w*SYT vs. *w*Mel, *w*SYT vs. *w*Au or *w*Mel vs. *w*Au), with a total of 2108-3202 SNPs and 234-457 indels (Table 2A, 3A). When classifying the SNPs based on the different genomic regions, it is clear that they are not randomly distributed in the genomes. We observed a strong overrepresentation of SNPs in the phage regions with approximately 40-50% of all SNPs located in these regions even though the phages represent only around 10%-15% of the genomes. This overrepresentation reflects the non-orthologous nature of several of the phage regions between the genomes (Figure S2). Additionally, we observed that there is a lower frequency of non-synonymous mutations in genes located in phage WO regions than in genes outside phage WO regions. The frequency of non-synonymous substitutions is only 35-40% in genes located in the phage WO regions, but around 65-69% in genes outside. Thus, the frequency of non-synonymous substitutions is much lower in genes from the phage WO regions compared to the estimated frequency of non-synonymous sites in single copy orthologs (76%). Such result indicates that selection might have acted during a longer time on the divergent and non-orthologous phage WO sequences (when they were present in other genomes) resulting in a lower ratio of non-synonymous to synonymous substitutions. Taken together, our analysis of SNPs suggests that the genes outside of the phage regions likely represent the “true” divergence between the close relatives, making the overall similarity between these genomes much higher than 99,8%.

Looking at our protein clusters, we found 19 clusters that were exclusive to the SYTMA genomes (Table S4). Twelve of them contain hypothetical proteins, one is an ANK protein, two are transposons, two are a putative toxin-antitoxin pair, one is a transcriptional regulator and the final protein is Geranyltranstransferase. Among the hypothetical proteins, we found the *w*Mel proteins WD0353 and WD0811, which were both seen to significantly affect the growth of yeast cells (Rice, et al. 2017), as well as the neighboring gene of WD0811, i.e. WD0812. However, none of the proteins in the 19 cluster were unique to this clade when compared to other *Wolbachia* genomes present in the nr database.

Among the SYTM genomes (excluding *w*Au) we identified five unique clusters (Table S4), of which two were unique to the SYTM clade, as we were unable to locate them in any other sequenced genome. Three of these proteins are located in the “Octomom” region in the *w*Mel genome (Chrostek, et al. 2013), analyzed in more detail below.

Finally, we identified 22 protein clusters that were unique to *w*SYT (Table S4). A majority of these (18) were located in two regions of the genome. One is a phage WO copy that is relatively highly diverged from the other genomes in our clustering (Figure S2) and the other is an unknown region with mostly hypothetical proteins that we call the “*Wolbachia Island*”, presented in more detail below.

##### The Octomom region

The Octomom region is made up of eight genes in *w*Mel and is known to be involved in overreplication and pathogenicity of the *w*MelPop strain (Chrostek and Teixeira 2015). It was previously noted as missing from the *w*Au genome (Iturbe-Ormaetxe, et al. 2005) and two of the proteins were shown to have been laterally transferred between *Wolbachia* and mosquitoes (Klasson, et al. 2009a; Woolfit, et al. 2009). In *w*SYT, the genes are directly flanking one of the phage WO regions (Figure 2A), similar to the homologous genes identified in the genome of the supergroup B strain from Culex mosquitoes *w*Pip (Klasson, et al. 2009a). In *w*Mel, the genes are located ca. 40 kbp away from the nearest phage WO region (Figure 2A).

**Figure 2.**
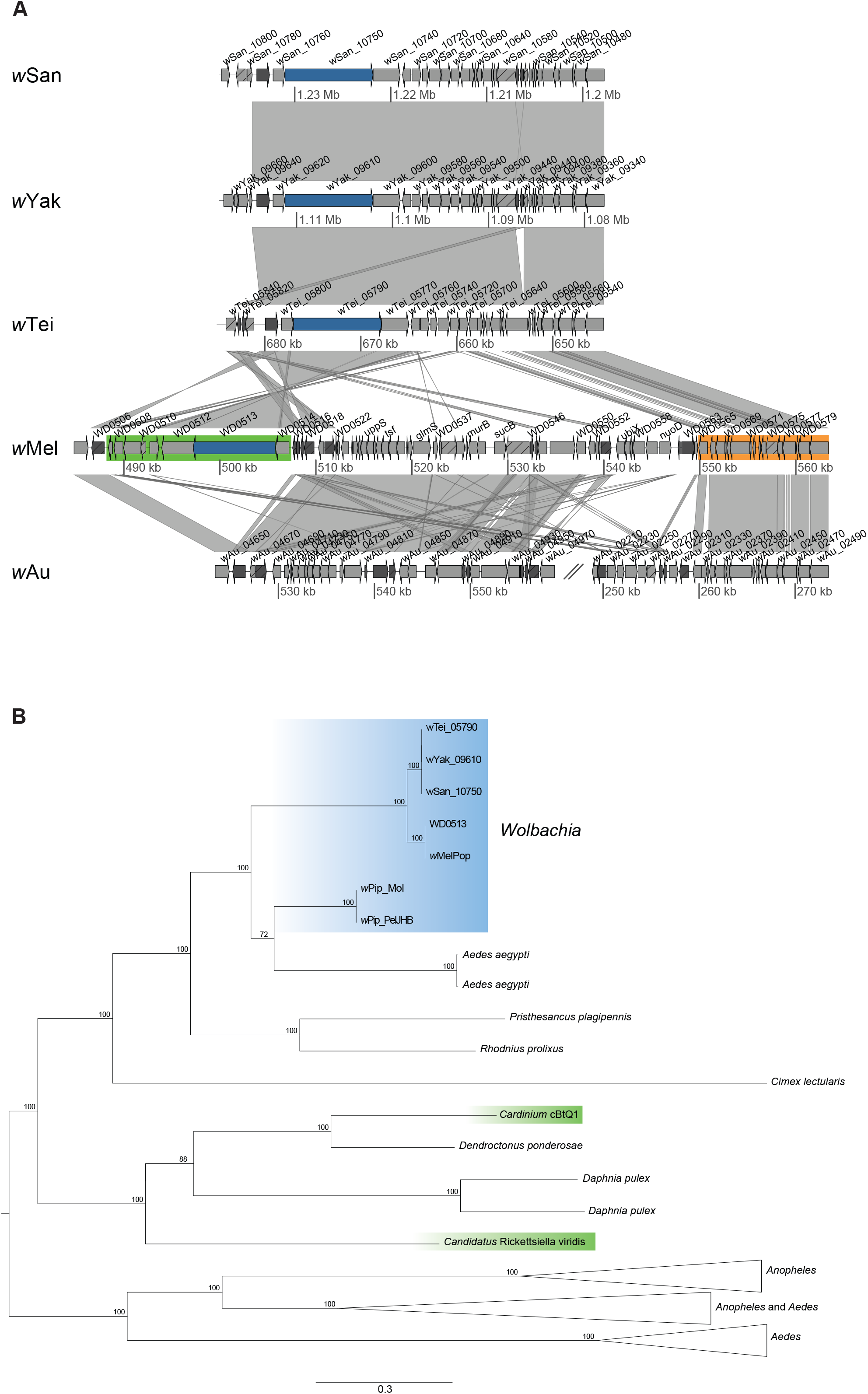
The Octomom region and neighboring genes. (**A**) Comparison of the Octomom region (highlighted in green in *w*Mel) in SYTMA genomes and neighboring phage-associate genes (highlighted in orange in *w*Mel). Putative functional genes are shown as grey arrows, while arrows with diagonal lines represent pseudogenes. Dark grey arrows are mobile elements. The blue arrow represents WD0513 and its homologs in *w*SYT. Similarity between sequences is indicated by the grey lines, where darker is more similar. (**B**) Maximum likelihood tree of the *w*Mel WD0513 gene and its homologs. An asterisk indicates 100% bootstrap support. Accession numbers for the proteins featured in the tree are available in Table S5.

Phylogenetic reconstruction of one of the proteins from the Octomom region (WD0513 in *w*Mel) shows that the *w*SYT proteins are most closely related to *w*Mel (Figure 2B), although the divergence of this gene between *w*SYT and *w*Mel is much higher than most other parts of the genomes (328-391 SNPs, ca. 15% of total SNPs and 11-17 indels, ca. 6% of total indels).

During this analysis, we also identified homologs of WD0513 in two other bacterial symbionts, the reproductive manipulator *Cardinium,* known to infect both insects and crustaceans (Edlund, et al. 2012; Schön, et al. 2019; Zchori-Fein and Perlman 2004), and the aphid endosymbiont “*Candidatus* Rickettsiella viridis” (Tsuchida, et al. 2010). WD0513 homologs were also found in several Hemiptera, including the predatory “assassin bug” *Pristhesancus plagipennis*, the bedbug *Cimex lectularis* and the kissing bug, *Rhodnius prolixus*, one of the vectors of Chagas disease. These new homologs branch outside of the *Wolbachia* clade and sit on long branches (Figure 2B), but are still closer to *Wolbachia* than any of the other symbionts. The only protein that goes inside the *Wolbachia* clade is one of the Salivary Gland Surface (SGS) genes of *Aedes aegypti* (Figure 2B), which was previously described (Klasson, et al. 2009a). The phylogenetic position of the WD0513 homologs from *Cardinium* and “*Candidatus* Rickettsiella viridis” suggests lateral transfers between these symbionts and their putative eukaryotic hosts.

##### The Wolbachia Island

The *Island* region in *w*SYT contains twelve genes and is flanked by a ca. 2.8 kbp repeat that includes a Group II intron encoded reverse transcriptase and a degraded transposase. Only one protein of the twelve, a putative addiction module toxin, clusters together with proteins from the other genomes. One additional protein contains a known protein domain, the C-terminal domain of DnaB-like helicase (PF03796). The remaining ten proteins have no hits to known protein domains or to any non-*Wolbachia* genome in nr.

Mainly five other *Wolbachia* genomes contain proteins or regions with significant similarity. The genomes of two B supergroup strains, *w*Con from the woodlouse *Cylisticus convexus* and *w*Lug from the planthopper *Nilaparvata lugens* both contain a region that is highly similar in content to the *w*SYT *Island* (Figure 3). Additionally, three other *Wolbachia* genomes, *w*Cle of supergroup F, *w*DacA of supergroup A and *w*Stri of supergroup B, have regions with significant similarity (Figure S3) but many of the proteins seem to have been pseudogenized in these genomes. Alternatively, some genes may appear fragmented due to the draft status of the genomes of *w*DacA and *w*Stri. The region is immediately flanked by genes with similarity to mobile elements on at least one side in all genomes (Figure 3), and one of the genes found in *w*Con and *w*Lug (pseudogenized in *w*SYT) has similarity with a phage portal protein.

**Figure 3.**
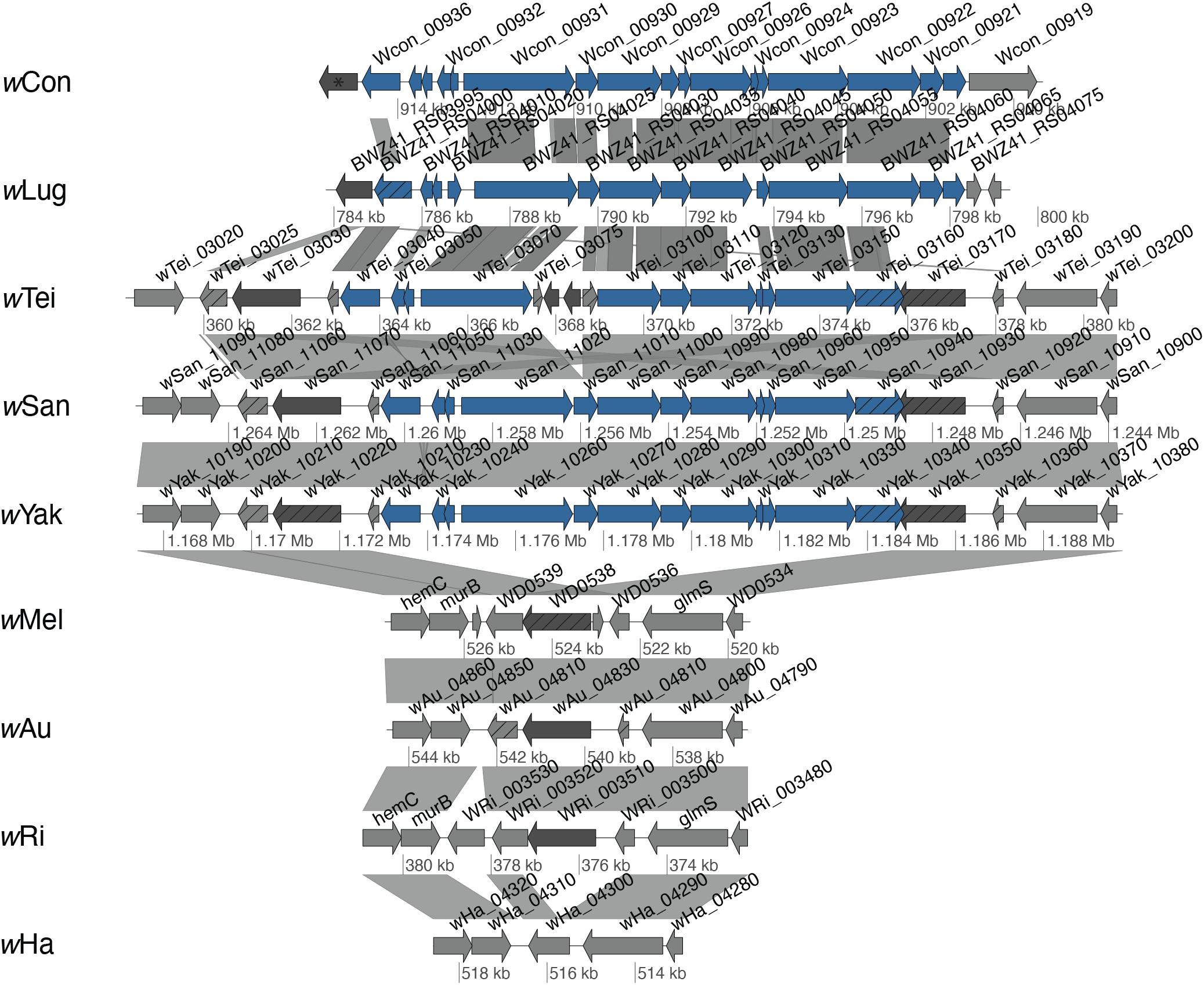
The *Wolbachia Island*. Comparison of the *Wolbachia Island* region in the genomes of *w*SYTMA, *w*Ri, *w*Ha, *w*Con and *w*Lug. Blue arrows show the Island genes in *w*SYT and their homologs in *w*Con and *w*Lug. Other putative functional genes are shown as grey arrows, while arrows with diagonal lines represent pseudogenes. Dark arrows represent mobile elements. Similarity between sequences is indicated by the grey lines, where darker is more similar.

The *Island* is completely missing in the genomes of the two very close wSYT relatives *w*Mel and *w*Au, as well as in the genomes of the two other supergroup A strains *w*Ri and *w*Ha in our analysis. However, the gene order flanking the region is conserved between all of them except one of the ends in *w*Tei. Hence, taken together, the most parsimonious explanation is that the *Island* has been inserted into the *w*SYT genomes through lateral gene transfer.

Interesting to note is that the plasmid pWCP, found in some *w*Pip strains, also contain putative toxin-antitoxin systems, a protein with the C-terminal domain of DnaB and a transposon (Reveillaud, et al. 2019). However, we did not find any other homologous proteins between the *Island* and the pWCP plasmid. We observe that in the draft genome of *w*Con the *Island* is located on a relatively small contig containing the same transposase on both ends. This suggests that the contig could represent an extrachromosomal circular DNA molecule such as a plasmid.

### Genetic variation associated with CI

#### The *cif* genes

In order to investigate how the *cif* genes correlate with CI phenotypes in our genomes, we searched for Cif proteins and performed phylogenetic reconstructions. We included the Cif proteins from the incomplete genomes of three additional strains with known CI phenotypes in *D. simulans* (*w*Ara, *w*Stv and *w*Triau) (Martinez, et al. 2015). Additionally, to get a good representation of the different Cif Types in the tree, Cif proteins from other complete *Wolbachia* genomes were included in the analysis.

We found CifB proteins in the genomes of all *mod^+^* strains in our analysis. They belonged to four of the different types (Types I-IV), with Type I proteins found in *w*SYT, *w*Mel and *w*Ha, Type II in *w*Ri, Type III in *w*No and Type IV in *w*Tei (Figure 4). Among the genomes of the *mod*^−^ strains, the *cifB* gene is completely absent from *w*Au and pseudogenized by a point mutation in *w*Ma.

**Figure 4.**
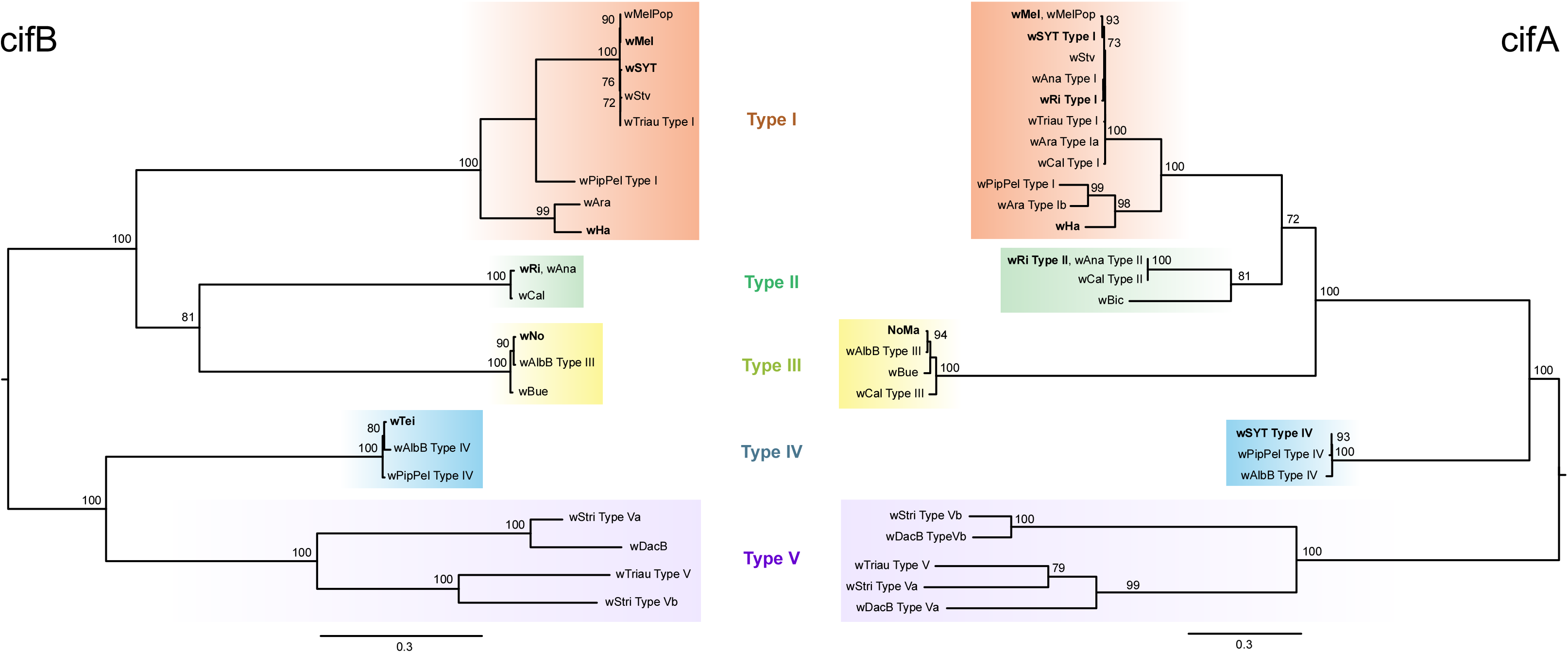
Phylogeny of the CI-associated genes *cifA-cifB* and their homologs. Maximum likelihood tree of (**A**) *cifB* and (**B**) *cifA* homologs. Five clusters named Type I-V are marked with different colours. Bootstrap supports below 70 are not shown.

CifA homologs were found in all strains except in the *resc^−^* strain *w*Au. The phylogenies of CifA and CifB were highly congruent (Figure 4) and genomes that contain a *mod* factor of one type also contain the *resc* factor of the same type, as previously observed (Lindsey, et al. 2018; Martinez, et al. 2020). Additional CifA proteins that did not have a corresponding CifB protein were also found in some genomes, for example in *w*SY, *w*Ma*, w*Ara and *w*Cal. Among our genomes, *w*Ma is the only strain that is unquestionably *mod^−^resc^+^*, as it can rescue the modification of *w*No but not itself induce CI. The presence of a CifA protein in *w*Ma that is identical to the CifA of *w*No thus makes sense based on previous observations.

##### The cif genes of wSYT

The very closely related *w*SYT strains differ in their CI properties, with *w*Tei inducing stronger CI and being able to rescue more strains than *w*SY. Hence, we expect that there are differences between Cif proteins in their genomes if these proteins are the main determinants of CI.

We observed that *w*Tei contains both a Type I and a Type IV CifB protein, while the *w*San and *w*Yak genomes only encode a Type I CifB protein (Figure 4B) plus two copies of a Type IV *cifB* gene that are both pseudogenized by frameshift mutations (Figure 5A). Thus, since the Type I CifB proteins are identical between the genomes and the Type IV CifB is probably non-functional in *w*SY, it is likely that the Type IV CifB protein in *w*Tei is the one that causes strong CI in *D. simulans*. Additionally, the Type I CifB proteins in *w*SYT are 112 amino acids shorter than the CifB protein of *w*Mel. The N-terminal truncation is due to an inversion (Figure 5B), also noted by Cooper, et al. (2019), that might have occurred via homologous recombination of a small inverted repeat (AGGACG). Although no functional domains are inferred in the truncated part of the protein, we might infer that the Type I CifB proteins of *w*SYT are either non-functional, based on the lack of CI induction by *w*SY seen by Martinez, et al. (2015), or not very effective in inducing CI, based on the results of Cooper, et al. (2017); Zabalou, et al. (2008).

**Figure 5.**
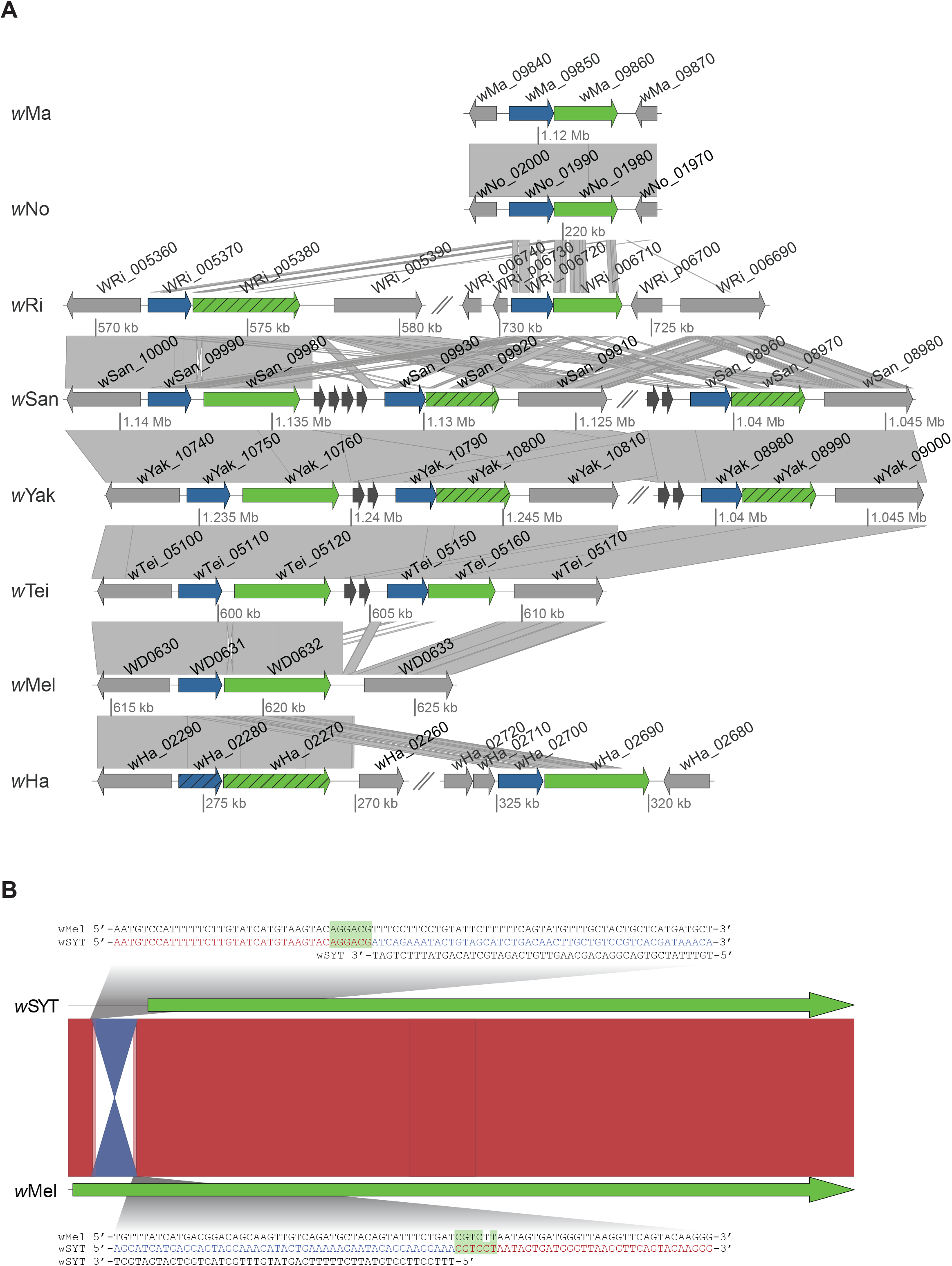
Comparison *cif* genes between genomes. (**A**) Comparison of *cifA*-*cifB* and homologs between the genomes used in this study. Grey arrows symbolize functional genes and arrows with diagonal lines indicate pseudogenes. *CifA-*homologs are blue and *cifB*-homologs are green. IS-elements are marked with darker grey. Similarity between sequences is indicated by the grey lines, where darker is more similar. **(B**) The inversion in the *w*SYT orthologs of the *cifB* gene are flanked by an inverted repeat shown in green. Similarity between sequences is indicated by red (forward match) or blue (reverse match) lines.

For CifA, each *w*SYT genome has one Type I protein plus either one Type IV protein (*w*Tei), or two identical Type IV proteins (*w*SY) (Figure 5A). The results from Zabalou et al. 2008 suggest that the rescue properties of *w*Tei and *w*SY are different, with *w*Tei being able to rescue the modification of *w*Mel while the *w*SY strains are not. Hence, even though these CifA proteins are very likely involved in rescue in *w*SYT, they alone can’t explain the differences in rescue potential between the *w*SYT genomes.

##### Origin and movement of cif genes in wSYT

The *cif* genes can move between unrelated *Wolbachia* genomes with the help of the WO phage, since most or all of the copies are located in phage regions and do not follow either strain or supergroup phylogenies (Figure 4). Using draft genomes, Cooper, et al. (2019) suggested that the Type IV *cifAB* genes of *w*SYT, might have been transferred laterally by the aid of flanking IS-elements (ISWpi1). Similar to Cooper, et al. (2019) we found that *w*Yak, as well as *w*Tei and *w*San, have ISWpi1 elements between the Type I and Type IV *cifAB* loci (Figure 6). However, when comparing the *w*SYT genomes, we did not find ISWpi1 elements in the same location at the other end. Additionally, the *w*SYT genomes are not syntenic through this entire phage region.

**Figure 6.**
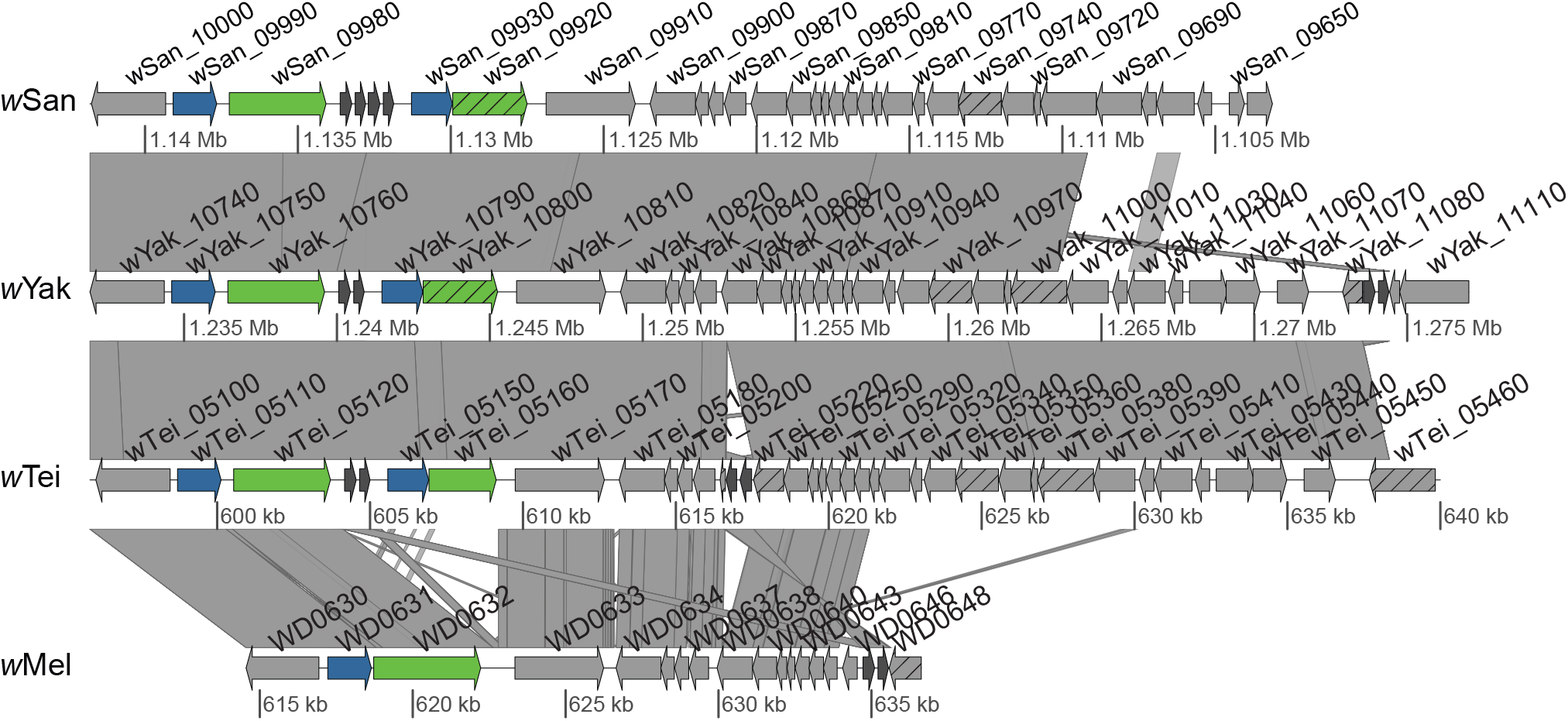
Comparison of the *cifA-cifB* region between SYTM genomes. *CifA-*homologs are marked in blue and *cifB*-homologs in green. Other putative functional genes are shown as grey arrows, while arrows with diagonal lines indicate pseudogenes. ISWpi1-elements are marked with a darker grey. Similarity between sequences is indicated by grey lines, where darker is more similar.

At least one duplication followed by several rearrangements of this region must have occurred in the *w*SY genomes, which makes it hard to infer a detailed scenario (Figure 6, Figure S4). However, we observe that the WO phage copy associated with the two Cif loci appears complete in *w*Tei, and that the WO phage connected to the Type IV *cifAB* locus in *w*Yak has the same content and gene order as *w*Tei (Figure S4) — even though it is found in a different genomic location. The same WO phage region also contains the Octomom WD0513 homolog at its other end, after which another ISWpi1 copy exists (Figure 6). Given that the WO phage found in connection with the Type IV *cifAB* genes appears complete in *w*Tei and *w*Yak, and that the proteins have a relatively consistent phylogenetic position throughout its full extent (Figure 4, Figure S4), it is most likely that the *cifAB* Type IV genes as well as the WD0513 homolog entered the *w*SYT genomes via a WO phage rather than via recombination of a DNA segment flanked by ISWpi1 elements. The close relationship between *w*SYT and *w*Pip for both the WD0513 homologs and Type IV CifAB proteins make this hypothesis highly plausible and suggests a supergroup B origin of the Type IV *cifAB* locus in *w*SYT. The presence of WD0513 and the Octomom region in *w*Mel might indicate that this WO phage was present in the ancestor of SYTM. In that case, the Type IV *cifAB* genes must have been lost in *w*Mel.

#### Other genes associated with *mod* and *resc*

To identify further gene candidates potentially associated with *mod* and *resc* functions, we compared patterns of gene absence/presence with the phenotypic expression of *mod* and *resc* and examined genes with significant divergence between strains that either induce or don’t induce CI.

##### Mod candidates

To identify proteins associated with modification, we looked for clusters of proteins that were present in all CI-inducing strains in our analysis but absent in the non-CI inducing strains. No protein clusters were identified by treating *w*SY as *mod^−^* together with *w*Au and *w*Ma. However, when treating *w*SY as *mod^+^* we identified two clusters (Table 4), one with the CifB proteins and one containing hypothetical proteins that are homologs of WD0462 in *w*Mel. The latter are found in one copy in all CI strains, but are pseudogenized in both *w*Ma and *w*Au (Figure 7). In several of the strains the protein contains the HAUS Augmin-like complex subunit 3, N-terminal domain (PF14932) (Figure 7). Interestingly, the *w*Mel protein WD0462 was seen to negatively affect growth when it was expressed in yeast under stress conditions (Rice, et al. 2017).

**Table 4:** Proteins associated with the CI phenotype.

| Protein | wMel locus tag |
| --- | --- |
| **Proteins associated with modification** |  |
| **Presence/Absence** |  |
| CifB | WD0632 |
| Hypothetical protein | WD0462 |
| **Divergence** |  |
| DNA directed RNA polymerase, beta/beta' subunits | WD0024 |
| NADH-quinone oxidoreductase subunit H | WD0159 |
| Acetyl/propionyl-CoA carboxylase, alpha subunit | WD0433 |
| **Genes with upstream variation in wSYT** |  |
| Ankyrin repeat domain protein | WD0292 |
| Hypothetical protein | WD0403 |
| S-adenosylmethionine synthase | WD0136 |
| Aspartate-semialdehyde dehydrogenase | WD0954 |
| **Proteins associated with rescue** |  |
| **Presence/Absence** |  |
| CifA | WD0631 |
| Phage replication protein RepA | WD0582,<br>WD0609 |
| Hypothetical protein | WD1187 |
| DNA recombination-mediator protein A | WD0092 |
| **Divergence** |  |
| M16 family peptidase | WD0762 |
| Folypolyglutamate synthase | WD1052 |

**Figure 7.**
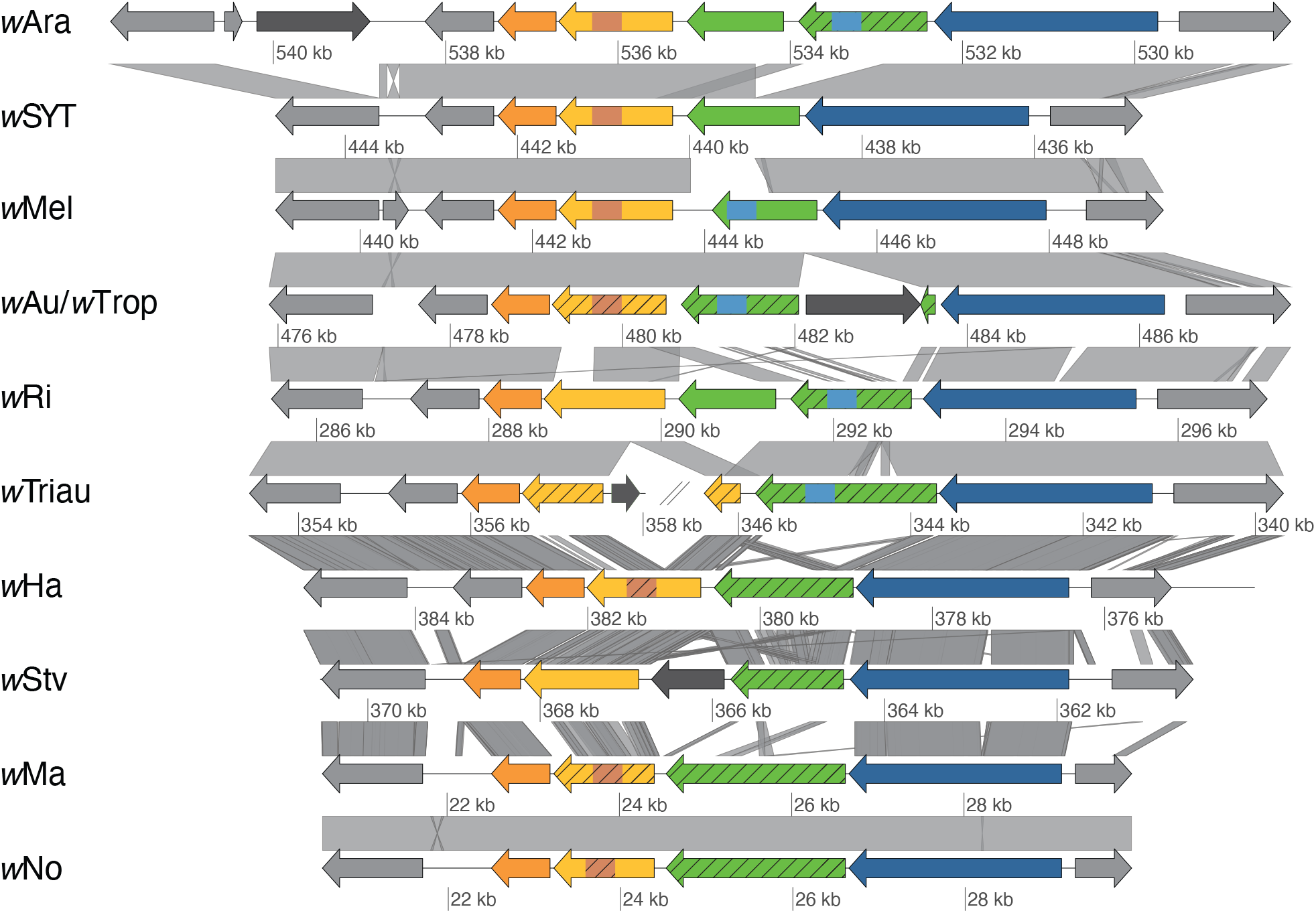
Comparison of the genomic region containing the *w*Mel gene WD0462. Yellow arrows represent WD0462 homologs, with light brown rectangles indicating predicted HAUS Augmin3 domain (PF14932). Green arrows represent WD0463 homologs, with light blue rectangles showing predicted AAA-ATPase domains (PF00004). Rectangles with a striped pattern indicate that the domain prediction was below the Pfam significance score. Orange and blue arrows show the genes *pyrF* and *valS* which flank the WD0462 and WD0463 loci in the analyzed genomes. Other putative functional genes are shown as grey arrows, while arrows with diagonal lines indicate pseudogenes. Dark arrows are mobile elements. Similarity between sequences is indicated by grey lines, where darker is more similar.

We further tested the link between WD0462 homologs and the CI phenotype by analyzing the status of this gene in the genome assemblies of the *Wolbachia* strains *w*Ara, *w*Stv, *w*Triau and *w*Tro, all of which have known CI phenotypes in *D. simulans* (Martinez, et al. 2015). Although these assemblies are incomplete, we see that our predictions are met in the *mod*^+^ strains *w*Ara and *w*Stv, which have functional WD0462 homologs, and in the *mod*^−^ *w*Tro, which has a pseudogenized copy. Only *w*Triau doesn’t follow our prediction, as this strain is *mod*^+^ but has a truncated and split WD0462 copy. However, we cannot exclude the possibility that a complete copy of the WD0462 may exist in the *w*Triau genome, or that the truncated gene is partially functional. It is worth noting that the neighboring gene WD0463 is a distant homolog of WD0462 that varies significantly between strains (Figure 7). In many of the strains, the protein homologs of WD0463 contain an AAA-ATPase domain (PF00004) (Figure 7).

To identify more genes potentially associated with *mod*, we searched for genes that are divergent between the CI and non-CI (or low CI for the *w*SY) genomes. We primarily considered genes that contained substitutions that separated the most closely related strains with different phenotypes, *w*SYT and *w*Ma/*w*No. Out of our 714 single copy protein clusters, we identified three genes that follow a pattern of divergence that could make them associated with *mod* i.e. they were identical between *w*San and *w*Yak but had at least one non-synonymous mutational difference with *w*Tei, and contained at least one non-synonymous mutational difference between *w*Ma and *w*No (Table 4). The three genes code for NADH-quinone oxidoreductase subunit H, Acetyl/propionyl-CoA carboxylase, alpha subunit and DNA directed RNA polymerase, beta/beta’ subunits. Based on parsimony, the mutations in NADH-quinone oxidoreductase subunit H occurred in *w*Ma and *w*Tei; in DNA directed RNA polymerase, beta/beta’ subunits (*rpoBC*) the mutations occurred in *w*No and *w*Tei; and in Acetyl/propionyl-CoA carboxylase, alpha subunit four mutations were exclusive to *w*No, one to *w*Ma and one to *w*SY. None of these mutations occur at the same positions in the gene in the different strains. Given the putative functions of these proteins, the very low divergence between strains with different phenotypes and the lack of parallel mutations, we believe that none of these genes are likely to be involved in *mod*. However, it is worth noting that the two CI-inducers, *w*Tei and *w*No, both have mutations in *rpoBC* and that the RpoBC protein was expressed in the ovaries of *Culex pipiens* (Buckeye) infected with a CI-inducing *Wolbachia* strain (LePage, et al. 2017).

We also used the *w*SYT genomes to look for mutations that might be involved in regulating expression levels by analyzing the upstream region of genes. We found mutations that differentiate *w*SY and *w*Tei in upstream regions of only six genes. Four of the genes are present in all CI-inducing strains and code for an ANK protein, a hypothetical protein, S-adenosylmethionine synthase and aspartate-semialdehyde dehydrogenase. Since non-coding regions are much less conserved, it is difficult to infer in which *Wolbachia* strains these mutations took place.

##### Resc candidates

To identify genes that are associated with rescue we looked for clusters that contained proteins from all genomes except *w*Au, since *w*Au is the only strain in our analysis that is unable to rescue the modification of any other strain. We identified four protein clusters that contained at least one protein from each genome except *w*Au (Table 4). Of these four clusters, two contain proteins that are located in phage WO and the remaining two are a hypothetical protein (WD1187 in *w*Mel) and DNA recombination-mediator protein A.

One of the proteins found in phage WO is CifA (WD0631 in *w*Mel) as described above. The other phage WO protein is the multifunctional phage replication protein RepA (Mardanov and Ravin 2006), previously annotated as Regulatory protein RepA. This protein is found in one copy in each supergroup B strain and in several copies in all supergroup A strains (WD0582 and WD0609 in *w*Mel) except *w*Au, which contains two copies with one frameshift each. Several of the other supergroup A genomes also contain additional pseudogenic copies of the gene (Figure S5). It is worth noting that a homolog of RepA was detected in the ovaries of *C. pipiens* (Buckeye) infected with a CI-inducing *Wolbachia* strain (LePage, et al. 2017).

The hypothetical protein WD1187 contains no conserved domains and is present in one copy in each *resc^+^* genome as well as a pseudogene with both a non-sense and a frameshift mutation in *w*Au (Figure S6). The protein has 3-4 transmembrane domains (3 with TMHMM and 4 with Phobius), and a very low similarity to the Endoplasmic-reticulum-associated protein degradation (ERAD)-associated E3 ubiquitin-protein ligase HRD1B from several plant species (*Brassica, Raphanus, Arabidopsis*) can be detected using two rounds of PSI-blast (14% identity over 70% of the protein, with E-values above 1). E3 ubiquitin ligases provide the specificity of ubiquitination by interacting with both the E2 ubiquitin-conjugating enzyme that carries the ubiquitin and the protein that is being ubiquitinated (Teixeira and Reed 2013). However, the protein is present in most *Wolbachia* genomes, including for example the mutualistic non-CI strain *w*Bm from the nematode *Brugia malayi*.

Finally, the DNA recombination-mediator protein A (previously DNA processing chain A - DprA) is found in one copy in all genomes except *w*Au, where it has a frameshift mutation (Figure S7).

As observed for CifA, all three additional rescue candidates were identical between the *w*SYT genomes. Hence, similarly to our *mod* analysis, we screened for divergent genes that correlate with *resc* properties, and identified two potential candidate genes under the assumption that the *resc* factor would have to be different between *w*SY and *w*Tei, *w*Au had to be different from all, and that this factor could be identical between *w*No and *w*Ma. The two genes code for a putative M16 family peptidase where one mutation seems to have occurred in *w*SY, one in *w*Tei and one in *w*Au; and for Folylpolyglutamate synthase where one mutation has occurred in *w*SY, one in *w*SYT and one in *w*Mel. Interestingly, an ortholog of the M16 family peptidase was found in the ovaries of *Culex pipiens* (Buckeye) infected with CI inducing *Wolbachia* strain (LePage, et al. 2017).

## DISCUSSION

*Wolbachia* participates in a remarkable variety of host phenotypes which range from mutualism to reproductive parasitism. Among these, CI stands out for its evolutionary implications to both host and symbiont as well as for the recent use in controlling insect-transmitted diseases. In spite of the scientific interest that CI creates, its genetic causes and mechanisms are still relatively poorly understood. This is partly due to the multiple host and symbiont factors that influence the phenotype. In the present study, we perform in-depth comparative genomic analyses of nine *Wolbachia* strains with known CI phenotypes in the *D. simulans* STC host background. By focusing on a single host background, we ensure that phenotypic variations between our strains are associated with symbiont rather than host factors. Using that strategy, we identify genetic features that are unique to specific strains, elucidate evolutionary patterns across closely related *Wolbachia,* and effectively pinpoint *Wolbachia* genes potentially associated with the *mod* and *resc* of CI.

### Rapid evolution of Wolbachia genomes and phenotypes are mediated by mobile elements

Phages and other mobile elements often occupy a relatively large proportion of *Wolbachia* genomes (Klasson, et al. 2009b; Wu, et al. 2004). They may also have a significant impact on *Wolbachia* ecology and evolution, since they frequently carry genes involved in host interaction and can be laterally transferred between strains (Bordenstein and Bordenstein 2016; Wang, et al. 2016). The phage WO associated *cif* genes, involved in CI, are prime examples of this phenomenon (LePage, et al. 2017; Madhav, et al. 2020; Martinez, et al. 2020).

Similar to previous studies (Ellegaard, et al. 2013; Gerth and Bleidorn 2016; Ishmael, et al. 2009), our comparisons show that phage WO regions contribute massively to the variation between closely related strains, as they contain a high proportion of the SNPs (SYTMA) as well as large gene content variability (all comparisons). Importantly, we observe that the Type IV *cif* genes of *w*Tei, likely causing the strong CI of this strain, are located in a phage WO region that was potentially transferred into *w*SYT from a Supergroup B donor. This *cif* pair may have been the only fully functional *cif* locus in the ancestor of *w*SYT, as the inversion in Type I *cifB* occurs in all three genomes while the pseudonization of Type IV *cifB* only occurs in *w*SY. Thus, the acquisition of this WO phage and consequently of the Type IV *cif* genes by the ancestor of *w*SYT may have had significant ecological importance for that *Wolbachia* lineage.

The same WO phage copy that carries the Type IV *cif* in *w*SYT is also associated with the Octomom gene WD0513, implicated in titer regulation of the *w*MelPop strain (Chrostek and Teixeira 2015; Duarte, et al. 2020). The location of WD0513 next to a phage in both *w*SYT and *w*Pip, as well as the sporadic presence of the Octomom region in *Wolbachia* genomes, suggest that the region is often laterally transferred by phage WO. Furthermore, it supports the claim that the Octomom region in *w*Mel was also originally part of a WO phage (Klasson, et al. 2009a). Our results show that homologs of WD0513 are present not only in *Wolbachia* but also in a variety of arthropod lineages and in two other endosymbionts of arthropods, *Rickettsiella* and *Cardinium.* This suggests lateral transfers not only occur between *Wolbachia* strains and *Wolbachia* and hosts, but potentially also between *Wolbachia* and other endosymbionts as well as their hosts. Although the mechanisms behind such transfers are unknown, phages or other mobile elements are likely involved. The WO phage is a likely culprit in *Wolbachia* transfers (Bordenstein and Bordenstein 2016), but less is known about mobile elements in “*Candidatus* Rickettsiella viridis” and *Cardinium*. We note, however, that the genome of “*Candidatus* Rickettsiella viridis” has one prophage region (Nikoh, et al. 2018) and that some *Cardinium* strains carry plasmids (Stouthamer, et al. 2019) that potentially could facilitate lateral transfers.

The novel “*Wolbachia Island*” also shows evidence of lateral transfer from Supergroup B into *w*SYT. We observed a few similarities between the types of genes found in the *Island* and those located on the pWCP plasmid of some *w*Pip strains (Reveillaud, et al. 2019). Although no direct conclusion can be made, we speculate that the *Island* could have been derived from an integrated plasmid. Since both plasmid- and phage-associated genes are often implicated in environment and host interaction in symbionts (Harumoto and Lemaitre 2018; Weldon, et al. 2013; Wernegreen and Moran 2001), the *Island* genes could thus potentially carry such functions.

Taken together, our observations suggest that mobile elements are drivers of rapid evolution in *Wolbachia*, where they mediate gain and loss of genes involved in ecologically important traits such as titer variation and CI.

### Factors associated with induction and rescue of CI

*Wolbachia* CI is a complex phenotype whose expression depends not only on the *cif* genes but also on a variety of factors including temperature (Bordenstein and Bordenstein 2011), *Wolbachia* titer and localization within the host (Breeuwer and Werren 1993; Clark, et al. 2003; Ikeda, et al. 2003; Veneti, et al. 2003), host species (Boyle, et al. 1993; McGraw, et al. 2001; Poinsot, et al. 1998; Zabalou, et al. 2008), host male age, mating rate and development time (Reynolds and Hoffmann 2002) as well as the age of the father and grandmother of the mating male (Awrahman, et al. 2014; Layton, et al. 2019). Here we take advantage of the reduced host-associated phenotypic variation in our dataset to generate new insight on how the *cif* genes are linked to inter-strain compatibility, identify novel gene candidates for the CI *mod* and *resc* functions, and to discuss how *Wolbachia* titer may be implicated in CI.

#### The *cif* genes

The *cif* genes are the main *Wolbachia* factors implicated in CI, with *cifB* linked to *mod* and *cifA* either to *resc* or both *mod* and *resc* (Beckmann, et al. 2019; Shropshire and Bordenstein 2019). These roles imply that a strain carrying a functional *cifB* also needs a functional *cifA* to be compatible with itself (Martinez, et al. 2020). We observe such a pattern in our strains, in which both *cifA* and *cifB* are intact or *cifB* is pseudogenized either alone or in combination with *cifA*. However, *cifA* is never pseudogenized alone. Additionally, the association of *cifA* with *resc* is supported by the fact that all of our *resc*^+^ strains have at least one putatively functional copy of *cifA*.

A similar association between *cifB* and *mod* implies that all *mod^+^* strains should carry a putatively functional copy of *cifB.* This is indeed the case for the strains *w*Ha, *w*Mel, *w*No, *w*Ri and *w*Tei. However, the *w*SY strains are also *mod^+^* according to Zabalou, et al. (2008) and Cooper, et al. (2017) but don’t have any fully intact *cifB* genes. We must then either consider that both strains don’t cause CI or that their truncated Type I CifB is at least partially functional. If the latter case is true, a weaker CifB function would also support recent findings that mutations outside of the main described domains of the Cif proteins can affect their CI properties (Shropshire, et al. 2020a). Reduced CifB functionality due to truncation could then, perhaps together with low infection titer (see discussion about titer below), be one of the reasons why *w*SY cause weaker CI in *D. simulans* in comparison to other strains that carry Type I CifB, such as *w*Mel and *w*Ha (Zabalou, et al. 2008).

The analysis of the *cif* genes in our genomes supports previous observations that strains carrying phylogenetically related *cif* tend to be compatible with each other (Bonneau, et al. 2018; Shropshire, et al. 2020b). Similarity between *cif* genes can explain why the *w*SYT strains can rescue each other’s modification, as the three strains have identical Type I and IV *cifA* genes, and why *w*Ma can rescue *w*No, as they have identical Type III *cifA* genes. However, several discrepancies remain regarding the observed patterns of *cif* genes in different strains and their published CI phenotypes in *D. simulans*. First, Zabalou et al. showed that the three *w*SYT strains can rescue *w*Ri, but according to our analysis none of the *w*SYT genomes possess a Type II CifA homolog, and Type II is the only complete CifB in the *w*Ri genome. Secondly, the NoMa strains were seen to partially rescue *w*Tei (Zabalou, et al. 2008) but their CifA homolog is of Type III rather than Type IV, which is the CifB type likely causing CI in *w*Tei. Additionally, we observed that all CI-inducing supergroup A genomes in our dataset contain copies of Type I CifA-B, but only *w*Mel and *w*Ha seem to maintain both Type I genes intact, whereas *w*SYT and *w*Ri only encode an intact CifA. Thus, based on CifA and CifB being the *resc* and *mod* factors, *w*SYT, *w*Ri and *w*Ha should be able to rescue *w*Mel. However, this is only partly in agreement with the results of Zabalou, et al. (2008), as *w*Ri and *w*Tei rescue the CI induced by *w*Mel, but *w*SY don’t. According to the same study, *w*SYT and *w*Ri cannot rescue the modification induced by *w*Ha even though *w*Ha only has an intact *cifB* of Type I and both *w*SYT and *w*Ri have intact *cifA* genes of Type I. In this case, it is worth noting that the Type I *cif* genes of *w*Ha are in a distinct subclade within the Type I phylogeny compared to those of *w*SYT and *w*Ri (Figure 4). Hence, further experiments are necessary to investigate whether the two subclades of Type I represent distinct Types in the sense that *cif* genes from one cannot rescue modifications caused by genes from the other.

#### Are there more *Wolbachia* genes involved in CI than *cif*?

Since the *cif* genes cannot explain all phenotypic variation in CI between our strains when they are in the same host background, we conclude that other genes must be involved in the phenotype. Our search for *Wolbachia* genes associated with *mod* and *resc* recovered, apart from the *cif* genes, a few novel CI-associated genes. Among these, WD0462 of *w*Mel is particularly promising for having a role in *mod*, as it has high sequence variability between genomes (Figure 7), negatively affects growth when expressed in yeast under stress conditions (Rice, et al. 2017) and has a Haus-Augmin3-like complex subunit 3, N-terminal domain (PF14932). This protein domain is present in the Dgt3 protein of *D. melanogaster,* where it binds to the gamma-Tubulin ring complex (gamma-TuRC) and is required for the accumulation of the gamma-TuRC to the mitotic spindle (Chen, et al. 2017). The density of microtubules in the mitotic spindle is reduced without Augmin, which can lead to perturbed chromosome alignment and mitotic progression (Goshima, et al. 2008; Uehara, et al. 2009). Additionally, Augmin contributes to the generation of astral microtubules during mitosis, which are essential for checkpoint satisfaction and chromosome segregation (Hayward, et al. 2014). Interestingly, the neighboring gene, WD0463, is also highly variable between strains. Notably, only strains encoding the WD0462 protein with a significant prediction for the Haus-Augmin3 domain encode an intact WD0463 protein. Such a pattern suggests possible coevolution between the two proteins. The AAA-ATPase domain (PF00004) found in several of the homologs of WD0463 is associated with a variety of cell functions including cell-cycle regulation. In spite of these interesting characteristics, we note that WD0462 is not variable between *w*SYT and so cannot explain differences between them regarding CI strength and compatibility with other strains (Zabalou, et al. 2008). Further investigation of the putative role of WD0462 in CI and whether or how its function is linked to that of the *cif* genes are needed.

Other *mod* candidates that were identified due to their sequence divergence between our strains seem less likely to have a role in CI. However, potential effects on gene expression caused by mutations in RpoBC of *w*Tei and *w*No could perhaps affect the occurrence or strength of CI in those strains (see discussion about titer below). The same might be true for mutations in the upstream region of certain genes in *w*Tei in comparison to *w*SY, although expressional data from each genome would be needed to confirm or reject these hypotheses.

Among the genes associated with *resc*, the multifunctional phage protein RepA could be of interest, since it has the potential to regulate phage copy number which in turn might affect *Wolbachia* titer (Bordenstein, et al. 2006). A putative *resc*-related role of RepA is also supported by its presence in the proteome data from ovaries of the mosquito *C. pipiens* infected with a CI-inducing *Wolbachia* (LePage, et al. 2017). Recently, RepA was also identified as a CI candidate by Scholz, et al. (2020), who observed that the protein was present in many *w*Mel and *w*Ri-like metagenomically-assembled genomes (MAGs) but absent in several *w*Au-like MAGs.

One of our other *resc*-related proteins, the *w*Mel protein WD1187, has low similarity to some E3 ubiquitin ligases from plants. This is interesting given that at least one Cif Type is a de-ubiquitinating enzyme able to cleave both Lysine-48 and Lysine-63 linked ubiquitin (Beckmann, et al. 2017). Additionally, the concentration of E3 ligase in the cell is possibly a way to control the localization and fate of ubiquitinated proteins (Li, et al. 2003), which might indicate that either the protein expression level or *Wolbachia* titer could be important if the *resc* phenotype occurs through such a mechanism.

The last *resc*-associated protein, DprA, is necessary for natural transformation in several bacterial species (Duffin and Barber 2016; Smeets, et al. 2000; Takata, et al. 2005) and acquisition of genes via the gene transfer agent in *Rhodobacter capsulatum* (Brimacombe, et al. 2014). It has been seen to bind single-stranded DNA and interact with the RecA protein, thereby assisting in recombination (Mortier-Barriere, et al. 2007). We note that although no ortholog of DprA was detected in the ovaries of *w*Pip-infected *C. pipiens* (Buckeye), RecA was (LePage, et al. 2017). However, based on the known functions of this protein, it is hard to speculate how it might be involved in the rescue of CI.

It is worth noting that none of the CI-associated genes presented here are among those that were previously shown not to cause CI when transgenically expressed in *Wolbachia* free *D. melanogaster* (Perlmutter, et al. 2020). It is also important to consider that the potential CI-associated effect of these genes may be indirect rather than a direct role in *mod* or *resc*. An example of this would be an effect on *Wolbachia* traits such as titer and localization which in turn influence CI.

#### Is *Wolbachia* titer important for *resc*?

The variable ability of *w*SYT to rescue the modification of *w*Mel in *D. simulans* cannot be explained by either the *cif* genes or by our new CI gene candidates, since these are all identical in the three strains. Hence, we propose that the difference in rescue between the strains could be due to a quantitative rather than qualitative variation in the rescue factor. At least two lines of evidence support this suggestion. The rescue function of Type I CifA in *D. melanogaster* was shown to be dependent on expression level (Shropshire, et al. 2018), and CI strength is correlated with bacterial titer in eggs (Martinez, et al. 2015). As strong CI is clearly not caused by high *Wolbachia* titers in the egg, since modification occurs in sperm, this observation indicates that high bacterial titers are needed in eggs of *Wolbachia* strains causing strong CI. A likely interpretation of this is that high levels of the *resc* factor are needed in order to rescue a strong CI. Thus, one possibility is that the difference in rescue between *w*SYT is due to the higher *Wolbachia* titer of *w*Tei compared to *w*SY in the eggs of *D. simulans* (Martinez, et al. 2015), where rescue occurs. The higher titer of *w*Tei would then result in enough CifA production to rescue the modification of *w*Mel, while the lower titer of *w*SY would not allow them to do the same.

However, although the titer of *w*Tei is higher than that of *w*San and *w*Yak in *D. simulans* eggs, it is still much lower than that of *w*Ri or *w*Mel (Martinez, et al. 2015; Veneti, et al. 2004). Hence, an alternative hypothesis could be that higher levels of *cifA* in *w*Tei might be obtained independently of titer variation, for example through increased expression. In this context, it is interesting to note that we found a non-synonymous mutation between *w*Tei and *w*SY in the gene *rpoBC,* which encodes the beta and beta’ subunits of RNA polymerase. Although we did not find any differences in the upstream regions of known CI genes in *w*SYT, it is possible that other forms of gene regulation exist. It is also interesting to note that *w*SY have two copies of the Type IV *cifA* genes, and this might thus compensate partly for the low titer. Transcriptomic or proteomic data could provide valuable information regarding the expression levels of rescue genes in different strains and whether these correlate with the phenotypic observations. Regardless of whether the quantitative effect is due to titer or expression, our reasoning leads to the testable hypothesis that the right amount of the *resc* factor as well as a good fit between *mod* and *resc* factors are both needed to rescue the modification of a strong CI inducer such as *w*Mel in *D. simulans*.

One possibility is that if the *mod* and *resc* factors fit perfectly together by having evolved under selection in the same genome, bacterial titer (or the amount of expressed *resc* factor) matters less than if *mod* and *resc* have a worse fit. With a less than perfect fit, perhaps rescue might only be possible if the *resc* factor is overexpressed compared to the *mod* factor with a perfect fit, a model of “force by numbers”. If correct, this model predicts that *Wolbachia* strains with a high titer (or overexpressed *resc* factor) could more easily rescue the modification of other strains. This could give such strains an ecological advantage, as they would be potentially better at invading populations that are already infected with other CI-causing *Wolbachia* strains. In contrast, low titer strains, in which drift has created a worse fit between the *resc* and *mod* factors, would have difficulty to infect new host species that are more permissive to CI than their current host, since more *resc* factor might be needed to rescue the CI induced by the strain itself. This hypothesis might explain how “suicide” strains that don’t fully rescue themselves, such as *w*Tei after transfer into *D. simulans*, can evolve under low CI conditions when there is low selection pressure on the *resc* function, like *w*Tei in its natural host.

### *Wolbachia* factors influencing non-reproductive phenotypes

Due to the early establishment of *D. simulans* as a permissive host for a multitude of *Wolbachia* strains, several investigations of non-reproductive phenotypes have been performed (Clark, et al. 2003; Martinez, et al. 2014; Martinez, et al. 2015; Toomey and Frydman 2014; Toomey, et al. 2013; Veneti, et al. 2004; Veneti, et al. 2003).

All of our strains except for *w*No and *w*Ri were used for investigating *Wolbachia*-associated host protection against two RNA viruses (FHV and DCV) as well as female fecundity and lifespan in the *D. simulans* STC background (Martinez, et al. 2014). All strains except for *w*San provide protection against at least one virus, and *w*Ma, *w*Yak and *w*Ha provide significant although variable protection against FHV but not DCV. There is also a strong correlation between the levels of protection *Wolbachia* elicits against the two viruses, so it is likely that the mechanism involved is the same for both. Both *w*Tei and *w*Ha significantly increase host fecundity, whereas *w*Mel significantly decrease it and *w*San, *w*Au and wYak reduced lifespan with varying levels of significance (Martinez, et al. 2015). Five of our nine strains (*w*No, *w*Ri, *w*Mel, *w*Yak, *w*Tei) were also used to investigate *Wolbachia* tropism in the germline stem cell niche (GSCN) during oogenesis and in the hub of testes during spermatogenesis (Toomey and Frydman 2014; Toomey, et al. 2013). Among these, *w*No is highly targeted for GSCN, while *w*Ri has intermediate tropism and *w*Tei, *w*Mel and *w*Yak have low tropism. *w*Mel is highly targeted for the hub, with *w*Yak and *w*Ri showing intermediate tropism while *w*Tei and *w*No have low.

Although the closely related *w*SYT strains have variable phenotypes in four of the five cases discussed above, none of the protein clusters from our analyses contain any of the three strains uniquely. Hence, differently from the CI phenotype, it is unlikely that any of these non-reproductive phenotypes occur through the action of proteins that are uniquely involved in those functions. This is perhaps not surprising, as these phenotypes are continuous rather than discrete and several of them correlate with *Wolbachia* titer in somatic tissues of *D. simulans* (Martinez, et al. 2015). Thus, titer may be a crucial factor for the expression of *Wolbachia-* induced non-reproductive phenotypes.

Three genetic properties of *Wolbachia* have so far been seen to affect its titer. These are the number of copies of the Octomom region, as seen in the *Wolbachia* strain *w*MelPop (Chrostek and Teixeira 2015; Duarte, et al. 2020), the expression level of the *Wolbachia* actin-localizing effector 1 (*WalE1* - WD0830), whose overexpression results in increased symbiont titer (Sheehan, et al. 2016), and the presence of lytic WO phages, as observed in the wasp *Nasonia vitripennis* (Bordenstein, et al. 2006). Interestingly, one of the few things that clearly differ between the *w*SYT genomes is the number of phage WO regions, with *w*San having the largest amount of prophage DNA in its genome followed by *w*Yak and then *w*Tei. Currently, we don’t know if the WO phages in the *w*SYT genomes are expressed as lytic phage particles and whether they indeed affect *Wolbachia* titer, but the correlation between titer and amount of phage WO regions in the genome is intriguing. However, we can’t discard the possibility that different *Wolbachia* strains are controlled by different mechanisms, which would make it more difficult to pinpoint the exact genetic component involved in each phenotype, especially when more divergent strains are compared.

Finally, the embryonic distribution of *Wolbachia* was tested in eight of our nine strains (all except *w*Ha) (Veneti, et al. 2004), although in this study *w*SYT were tested in their natural hosts. While *w*Ri has a global distribution in the embryo, the SYTMA strains have a posterior distribution and NoMa an anterior. In this case we can’t take advantage of our closely related genomes, but it might still be interesting to investigate the 19 protein clusters specifically found in the SYTMA clade in order to elucidate if any of the proteins could be involved in *Wolbachia* posterior localization. Notably, two of the proteins were shown to have an effect on growth when expressed in yeast, and might thus be *Wolbachia* effectors (Rice, et al. 2017).

### *D. simulans* as a permissive host species

Several lines of evidence indicate that *D. simulans* is an unusually permissive host species for *Wolbachia*, including the multiple natural infections and the relatively easy transfection of new *Wolbachia* strains into this host (Merçot and Charlat 2004). This permissiveness of *D. simulans* to *Wolbachia* infections has contributed to CI studies and allowed us to detect *Wolbachia* phenotypic traits that could otherwise have remained unnoticed, as they are not as prominent in their natural hosts. However, it remains unclear why *D. simulans* is so susceptible to *Wolbachia* infections, since it is unknown what host proteins might interact with and control *Wolbachia* infection. The ability to carry and vertically transmit *Wolbachia* could come from a balance between limiting the infection, i.e. preventing systemic infections which might be lethal to the host, but still allowing some proliferation. One possibility is thus that the defenses of *D. simulans* are less effective against *Wolbachia* in general, so that an infection can be relatively easily established, but that they are still effective enough to prevent toxicity.

Additionally, CI seems to have a higher penetrance in *D. simulans* compared to some other related *Drosophila* species. One hypothesis for this observation is that the *D. simulans* lineage might have evolved to a large extent in the absence of any *Wolbachia* interaction or possibly only with *mod^−^* strains such as *w*Ma and *w*Au. This situation could thus have prevented the evolution of resistance to or alleviation of the *Wolbachia* CI mechanisms.

Further studies are needed to get to the root of why *D. simulans* can harbor such a plethora of *Wolbachia* infections and why this species might be particularly sensitive to the mechanism of CI.

## CONCLUSIONS

In the present study, we use complete genomes of closely related *Wolbachia* combined with their phenotypic data in the *D. simulans* STC host background to investigate *Wolbachia* evolution and genetic determinants of CI. Our analysis shows that transferring *Wolbachia* strains from other *Drosophila* into *D. simulans* doesn’t seem to create significant evolutionary pressures on any particular symbiont function, which corroborates this strategy for studying *Wolbachia.*

From an evolutionary perspective, we find support for phages and mobile elements playing an important role in *Wolbachia* ecology and evolution through lateral transfers of genes implicated in phenotype expression and host interaction. We find evidence that phylogenetically related *cif* genes tend to be compatible and that other genes apart from *cif* are associated with modification and rescue of CI among our genomes. Both the toxin-antidote and host-modification models of CI can be reconciliated with our observation, with the novel CI-associated genes potentially either affecting the affinity between CifA and CifB (TA) or influencing the host interactions that lead to modification and rescue (HM). Based on our results, we also speculate that *Wolbachia* titer could be the missing factor that explains variability in CI rescue capabilities of the *w*SYT strains as well as in other systems. A higher symbiont titer in eggs should favor rescue regardless of CI model, as it increases the probability that *resc*-associated proteins find their targets. High titer *Wolbachia* might thus be better at invading new host populations that already carry a CI-inducing *Wolbachia* strain. If true, infection titer is a highly relevant parameter to consider when designing strategies for using CI *Wolbachia* in biological control programs. Overall, we show that comparative genomics combined with phenotypic information is a powerful tool for studying *Wolbachia* evolution and identifying genes associated with various traits.

## MATERIALS AND METHODS

### DNA preparation and sequencing

All DNA samples used for Illumina and PacBio sequencing were produced using the protocol described in (Ellegaard, et al. 2013). Briefly, *Drosophila* flies were transferred to apple juice agar plates and allowed to oviposit for 2 hours, after which the eggs were collected, washed and dechorionated in 50% bleach and manually homogenized using a plastic pestle. Following centrifugation and filtration of the homogenate to enrich for *Wolbachia* cells, the resulting cell pellet was subjected to whole genome amplification using the Repli-g® midi kit (Qiagen), after which the DNA was purified using QIAamp® DNA mini kit (Qiagen) according to the manufacturer’s recommendations. Standard 350 bp fragment TruSeq libraries were constructed from DNA samples of 11 different *Wolbachia* strains (Table S1). All libraries were indexed and run together in one lane on an Illumina HiSeq 2500 machine, generating 2×100 bp sequence reads. Illumina libraries were produced and sequencing was performed at the SNP and SEQ platform, Uppsala. DNA from each of the five *Wolbachia* strains yielding complete genomes (Table S1) was used to create 5 kb fragment SMRTbell libraries. Each library was run using P6-C4 chemistry in one SMRT cell on the RSII PacBio instrument. PacBio libraries were produced and sequencing was performed at the Uppsala Genome Center, Uppsala.

### Genome assembly

Illumina reads were quality and adapter trimmed by Trimmomatic-0.22 (Bolger, et al. 2014) and error corrected based on k-mer frequencies using BayesHammer in SPAdes (Bankevich, et al. 2012). Corrected Illumina reads were assembled into contigs by AbySS (Simpson, et al. 2009), SPAdes (Bankevich, et al. 2012), IDBA (Peng, et al. 2012) and Velvet (Zerbino and Birney 2008) with k-mer sizes from 63 to 95 with an interval of four. Assembly statistics were calculated using the Perl script assemblathon_stats.pl (https://github.com/ucdavis-bioinformatics/assemblathon2-analysis). The N50 value, predicted genome size, total number of contigs and length of contigs were considered while selecting the best assembly from each assembler. In order to complete the draft genome, PacBio reads were obtained and assembled both independently using HGAP (Chin, et al. 2013) and together with Illumina reads using Spades. The best assembly was chosen based on the same criteria used for Illumina assemblies, and thereafter overlapping contigs were merged in Consed (Gordon, et al. 1998). Illumina and PacBio reads were mapped against the assemblies using BWA-mem (Li and Durbin 2009) with default parameters for Illumina reads and using PacBio settings for PacBio reads to check the correctness of the genome assembly. Gaps and inconsistency between the assemblies and the data were tested using PCR and resolved by direct Sanger sequencing of the PCR products. In cases where repeats were too large to span with PCR products, PacBio data was extensively inspected for consistency with the assembly and PCR products that go from unique to repeat sequence were also generated in most cases. All assemblies and read data were combined and curated using Consed.

### Annotation

The genomes were annotated using an automated annotation pipeline DIYA (Stewart, et al. 2009), as described in (Ellegaard, et al. 2013). Prodigal (Hyatt, et al. 2010) was used to predict the protein coding genes, while GenePRIMP (Pati, et al. 2010) and blastx were used to identify pseudogenes. tRNAscan-SE (Lowe and Eddy 1997) and RNAmmer (Lagesen, et al. 2007) were used to predict tRNA and rRNA, respectively. All predicted proteins were searched against the UniProt database and previously annotated *Wolbachia* proteomes using blastp. PFAM domains were identified using pfam_scan.pl (Li, et al. 2015). Mummer was used to predict repeats (Kurtz, et al. 2004). After automated annotation, all data was collected in Artemis (Rutherford, et al. 2000) and used to manually curate each genome. Repeats were annotated using by nucmer from the Mummer 3 package (Kurtz, et al. 2004) with a minimum of 300 bp and 95% identity. Phage regions were annotated manually by comparing the gene content to previously published *Wolbachia* genomes.

### Variant calling

Quality and adapter trimmed Illumina reads of transinfected (for *w*San, *w*Yak and *w*Tei) or introgressed (*w*Au) *Wolbachia* strains were aligned against each of the other finished genomes as well as against their respective genome with BWA-mem. *Wolbachia* strains from their natural hosts were also aligned against their respective genome with BWA-mem and subsequently sorted and marked for duplicates with the Picard toolkit (http://broadinstitute.github.io/picard/). For each set of aligned reads, indels were realigned with the IndelRealigner from GATK (McKenna, et al. 2010) and SNPs and Indels were called using the Haplotypecaller from GATK with a ploidy of 1. SNPs and Indels were filtered separately using the GATK best practice settings but removing the criteria for haplotype score and increasing the QD threshold to 15. To avoid spurious variant calls, only sites with a minimum read depth of 10 were used. snpEff (Cingolani, et al. 2012) was used to create a specific database for each of the five complete and annotated genomes and to identify the effect and location of each variant within them.

### Clustering and phylogenetic analyses

The proteomes of the nine *Wolbachia* strains were clustered using OrthoMCL (Abascal and Valencia 2003) with an inflation value of 1.5. For genes found in clusters containing a single copy in each genome, nucleotide sequences were extracted, translated to proteins, aligned using mafft-linsi (Katoh and Standley 2013) and backtranslated to nucleotides. Phylogenetic trees were constructed for each of these genes individually and for a concatenated alignment of all of them using RAxML Version 8.1.16 (Stamatakis 2006) with the GTRGAMMA model and 100 rapid bootstraps. The same procedure was used for all gene alignments analyzed. Synonymous and non-synonymous substitution rates were calculated using codeml from the PAML package (Yang 2007).

For the phylogenetic analysis of WD0513, all non-identical *Wolbachia* homologs found in our clusters were searched against the nr database using blastp. Proteins in the database that covered at least 80% of the length of the query and had an e-value smaller than e^−05^ were aligned to the homologs from the nine *Wolbachia* genomes using mafft-linsi and trimmed using trimAl (Capella-Gutierrez, et al. 2009) with the –automated1 setting. Phylogenies were inferred using RAxML Version 8.1.16 with the PROTGAMMAAUTO model and 100 rapid bootstraps. In a majority of cases, LG was the best scoring amino acid model that was used to create the phylogenetic tree. All trees were visualized in FigTree (http://tree.bio.ed.ac.uk/software/figtree/). For the phylogenetic analysis of the *cif* genes, CifA and CifB proteins from our genomes were combined with representative Cif proteins of Type I-V chosen among those featured in Martinez, et al. (2020). The resulting protein set was aligned, trimmed and used for phylogenetic reconstructions with RAxML as described above for WD0513. Domains prediction for WD462 and WD463 were made with the online implementation of pfamscan (https://www.ebi.ac.uk/Tools/pfa/pfamscan/) using default values.

All gene comparison figures were made using genoPlotR (Guy, et al. 2010), after blasting each genome against every other genome using blastn for close relatives and blastp for distant relatives.

## Acknowledgements

The authors thank Roel van Eijk for technical assistance running PCRs, Lina Juzokaite for DNA extractions for PacBio sequencing, and Kostas Bourtzis for providing fly lines as well as contributing with advice and support throughout. Illumina sequencing was performed at the SNP&SEQ Technology Platform and PacBio sequencing was performed at Uppsala Genomic Center in Uppsala, Sweden, both of which are part of the Swedish National Genomics Infrastructure. This work was supported by grants from Formas (2009-378) and The Swedish research council VR (2014-4353) to LK.

## Author contributions

LK conceived and designed the study. JJ, LK and MG performed assemblies. JJ, GCB, LK and MG performed annotations. LK, GCB and JJ performed comparative and phylogenetic analyses. MG performed DNA extractions. LK and GCB wrote the paper with contribution from JJ. All authors read and approved the manuscript.

## Data availability

The data underlying this article are available at NCBI and can be accessed from the BioProject PRJNA694504. All individual accession numbers can be found in Supplementary Table 1.

## References

Abascal, F, Valencia, A 2003. Automatic annotation of protein function based on family identification. Proteins 53: 683–692. doi: 10.1002/prot.10449

Awrahman, ZA, Champion de Crespigny, F, Wedell, N 2014. The impact of *Wolbachia*, male age and mating history on cytoplasmic incompatibility and sperm transfer in *Drosophila simulans*. Journal of Evolutionary Biology 27: 1–10. doi: 10.1111/jeb.12270

Bankevich, A, Nurk, S, Antipov, D, Gurevich, AA, Dvorkin, M, Kulikov, AS, Lesin, VM, Nikolenko, SI, Pham, S, Prjibelski, AD, et al. 2012. SPAdes: a new genome assembly algorithm and its applications to single-cell sequencing. Journal of Computational Biology 19: 455–477. doi: 10.1089/cmb.2012.0021

Beckmann, JF, Bonneau, M, Chen, H, Hochstrasser, M, Poinsot, D, Merçot, H, Weill, M, Sicard, M, Charlat, S 2019. The toxin-antidote model of cytoplasmic Incompatibility: genetics and evolutionary implications. Trends in Genetics 35: 175–185. doi: 10.1016/j.tig.2018.12.004

Beckmann, JF, Ronau, JA, Hochstrasser, M 2017. A *Wolbachia* deubiquitylating enzyme induces cytoplasmic incompatibility. Nat Microbiol 2: 17007. doi: 10.1038/nmicrobiol.2017.7

Bolger, AM, Lohse, M, Usadel, B 2014. Trimmomatic: a flexible trimmer for Illumina sequence data. Bioinformatics 30: 2114–2120. doi: 10.1093/bioinformatics/btu170

Bonneau, M, Atyame, C, Beji, M, Justy, F, Cohen-Gonsaud, M, Sicard, M, Weill, M 2018. *Culex pipiens* crossing type diversity is governed by an amplified and polymorphic operon of *Wolbachia*. Nat Commun 9: 319. doi: 10.1038/s41467-017-02749-w

Bordenstein, SR, Bordenstein, SR 2016. Eukaryotic association module in phage WO genomes from *Wolbachia*. Nat Commun 7: 13155. doi: 10.1038/ncomms13155

Bordenstein, SR, Bordenstein, SR 2011. Temperature affects the tripartite interactions between bacteriophage WO, *Wolbachia*, and cytoplasmic incompatibility. PLoS ONE 6: e29106. doi: 10.1371/journal.pone.0029106

Bordenstein, SR, Marshall, ML, Fry, AJ, Kim, U, Wernegreen, JJ 2006. The tripartite associations between bacteriophage, *Wolbachia*, and arthropods. PLOS Pathogens 2: e43. doi: 10.1371/journal.ppat.0020043

Bordenstein, SR, O’Hara, FP, Werren, JH 2001. *Wolbachia*-induced incompatibility precedes other hybrid incompatibilities in *Nasonia*. Nature 409: 707–710. doi: 10.1038/35055543

Bossan, B, Koehncke, A, Hammerstein, P 2011. A New Model and Method for Understanding *Wolbachia*-Induced Cytoplasmic Incompatibility. PLoS ONE 6: e19757. doi: 10.1371/journal.pone.0019757

Boyle, L, O’Neill, SL, Robertson, HM, Karr, TL 1993. Interspecific and intraspecific horizontal transfer of *Wolbachia* in *Drosophila*. Science 260: 1796–1799.

Breeuwer, JA, Werren, JH 1993. Cytoplasmic incompatibility and bacterial density in *Nasonia vitripennis*. Genetics 135: 565–574.

Brimacombe, CA, Ding, H, Beatty, JT 2014. *Rhodobacter capsulatus* DprA is essential for RecA-mediated gene transfer agent (RcGTA) recipient capability regulated by quorum-sensing and the CtrA response regulator. Molecular Microbiology 92: 1260–1278. doi: 10.1111/mmi.12628

Capella-Gutierrez, S, Silla-Martinez, JM, Gabaldon, T 2009. trimAl: a tool for automated alignment trimming in large-scale phylogenetic analyses. Bioinformatics 25: 1972–1973. doi: 10.1093/bioinformatics/btp348

Charlat, S, Riegler, M, Baures, I, Poinsot, D, Stauffer, C, Mercot, H 2004. Incipient evolution of *Wolbachia* compatibility types. Evolution 58: 1901–1908.

Chen, H, Ronau, JA, Beckmann, JF, Hochstrasser, M 2019. A *Wolbachia* nuclease and its binding partner provide a distinct mechanism for cytoplasmic incompatibility. Proceedings of the National Academy of Sciences 116: 22314. doi: 10.1073/pnas.1914571116

Chen, JWC, Chen, ZA, Rogala, KB, Metz, J, Deane, CM, Rappsilber, J, Wakefield, JG 2017. Cross-linking mass spectrometry identifies new interfaces of Augmin required to localise the γ-tubulin ring complex to the mitotic spindle. Biology Open 6: 654. doi: 10.1242/bio.022905

Chin, CS, Alexander, DH, Marks, P, Klammer, AA, Drake, J, Heiner, C, Clum, A, Copeland, A, Huddleston, J, Eichler, EE, et al. 2013. Nonhybrid, finished microbial genome assemblies from long-read SMRT sequencing data. Nature Methods 10: 563–569. doi: 10.1038/nmeth.2474

Chrostek, E, Marialva, MSP, Esteves, SS, Weinert, LA, Martinez, J, Jiggins, FM, Teixeira, L 2013. *Wolbachia* Variants Induce Differential Protection to Viruses in *Drosophila melanogaster:* A Phenotypic and Phylogenomic Analysis. PLoS Genetics 9: e1003896. doi: 10.1371/journal.pgen.1003896

Chrostek, E, Teixeira, L 2015. Mutualism breakdown by amplification of *Wolbachia* genes. PLoS Biol 13: e1002065. doi: 10.1371/journal.pbio.1002065

Cingolani, P, Platts, A, Wang le, L, Coon, M, Nguyen, T, Wang, L, Land, SJ, Lu, X, Ruden, DM 2012. A program for annotating and predicting the effects of single nucleotide polymorphisms, SnpEff: SNPs in the genome of *Drosophila melanogaster* strain w1118; iso-2; iso-3. Fly 6: 80–92. doi: 10.4161/fly.19695

Clark, ME, Veneti, Z, Bourtzis, K, Karr, TL 2003. *Wolbachia* distribution and cytoplasmic incompatibility during sperm development: the cyst as the basic cellular unit of CI expression. Mechanisms of Development 120: 185–198. doi: S0925477302004240 [pii]

Cooper, BS, Ginsberg, PS, Turelli, M, Matute, DR 2017. *Wolbachia* in the *Drosophila yakuba* Complex: Pervasive Frequency Variation and Weak Cytoplasmic Incompatibility, but No Apparent Effect on Reproductive Isolation. Genetics 205: 333–351. doi: 10.1534/genetics.116.196238

Cooper, BS, Vanderpool, D, Conner, WR, Matute, DR, Turelli, M 2019. *Wolbachia* Acquisition by *Drosophila yakuba*-Clade Hosts and Transfer of Incompatibility Loci Between Distantly Related *Wolbachia*. Genetics 212: 1399. doi: 10.1534/genetics.119.302349

Duarte, EH, Carvalho, A, López-Madrigal, S, Teixeira, L 2020. Regulation of *Wolbachia* proliferation by the amplification and deletion of an addictive genomic island. bioRxiv: 2020.2009.2008.288217. doi: 10.1101/2020.09.08.288217

Duffin, PM, Barber, DA 2016. DprA is required for natural transformation and affects pilin variation in *Neisseria gonorrhoeae*. Microbiology 162: 1620–1628. doi: 10.1099/mic.0.000343

Edlund, A, Ek, K, Breitholtz, M, Gorokhova, E 2012. Antibiotic-Induced Change of Bacterial Communities Associated with the Copepod *Nitocra spinipes*. PLoS ONE 7: e33107. doi: 10.1371/journal.pone.0033107

Ellegaard, KM, Klasson, L, Naslund, K, Bourtzis, K, Andersson, SG 2013. Comparative genomics of *Wolbachia* and the bacterial species concept. PLoS Genetics 9: e1003381. doi: 10.1371/journal.pgen.1003381

Fast, EM, Toomey, ME, Panaram, K, Desjardins, D, Kolaczyk, ED, Frydman, HM 2011. *Wolbachia* enhance *Drosophila* stem cell proliferation and target the germline stem cell niche. Science 334: 990–992. doi: 10.1126/science.1209609

Flores, HA, O’Neill, SL 2018. Controlling vector-borne diseases by releasing modified mosquitoes. Nature Reviews Microbiology 16: 508–518. doi: 10.1038/s41579-018-0025-0

Gerth, M, Bleidorn, C 2016. Comparative genomics provides a timeframe for *Wolbachia* evolution and exposes a recent biotin synthesis operon transfer. Nat Microbiol 2: 16241. doi: 10.1038/nmicrobiol.2016.241

Gordon, D, Abajian, C, Green, P 1998. Consed: a graphical tool for sequence finishing. Genome Research 8: 195–202.

Goshima, G, Mayer, M, Zhang, N, Stuurman, N, Vale, RD 2008. Augmin: a protein complex required for centrosome-independent microtubule generation within the spindle. The Journal of cell biology 181: 421–429. doi: 10.1083/jcb.200711053

Guy, L, Kultima, JR, Andersson, SG 2010. genoPlotR: comparative gene and genome visualization in R. Bioinformatics 26: 2334–2335. doi: 10.1093/bioinformatics/btq413

Harumoto, T, Lemaitre, B 2018. Male-killing toxin in a bacterial symbiont of *Drosophila*. Nature 557: 252–255. doi: 10.1038/s41586-018-0086-2

Hayward, D, Metz, J, Pellacani, C, Wakefield, JG 2014. Synergy between multiple microtubule-generating pathways confers robustness to centrosome-driven mitotic spindle formation. Developmental Cell 28: 81–93. doi: 10.1016/j.devcel.2013.12.001

Hedges, LM, Brownlie, JC, O’Neill, SL, Johnson, KN 2008. *Wolbachia* and virus protection in insects. Science 322: 702. doi: 322/5902/702 [pii] 10.1126/science.1162418

Hurst, LD 1991. The evolution of cytoplasmic incompatibility or when spite can be successful. Journal of Theoretical Biology 148: 269–277. doi: https://doi.org/10.1016/S0022-5193(05)80344-3

Hyatt, D, Chen, G-L, Locascio, PF, Land, ML, Larimer, FW, Hauser, LJ 2010. Prodigal: prokaryotic gene recognition and translation initiation site identification. BMC Bioinformatics 11: 119. doi: 10.1186/1471-2105-11-119

Ikeda, T, Ishikawa, H, Sasaki, T 2003. Infection density of *Wolbachia* and level of cytoplasmic incompatibility in the Mediterranean flour moth, *Ephestia kuehniella*. Journal of Invertebrate Pathology 84: 1–5. doi: S002220110300106X [pii]

Ishmael, N, Dunning Hotopp, JC, Ioannidis, P, Biber, S, Sakamoto, J, Siozios, S, Nene, V, Werren, J, Bourtzis, K, Bordenstein, SR, et al. 2009. Extensive Genomic Diversity of Closely Related *Wolbachia* Strains. Microbiology. doi: mic.0.027581-0 [pii] 10.1099/mic.0.027581-0

Iturbe-Ormaetxe, I, Burke, GR, Riegler, M, O’Neill, SL 2005. Distribution, expression, and motif variability of ankyrin domain genes in *Wolbachia* pipientis. Journal of Bacteriology 187: 5136–5145. doi: 187/15/5136-a [pii] 10.1128/JB.187.15.5136-5145.2005

Jaenike, J 2007. Spontaneous emergence of a new *Wolbachia* phenotype. Evolution 61: 2244–2252.

Katoh, K, Standley, DM 2013. MAFFT multiple sequence alignment software version 7: improvements in performance and usability. Molecular Biology and Evolution 30: 772–780. doi: 10.1093/molbev/mst010

Klasson, L, Kambris, Z, Cook, PE, Walker, T, Sinkins, SP 2009a. Horizontal gene transfer between *Wolbachia* and the mosquito *Aedes aegypti*. BMC Genomics 10: 33. doi: 10.1186/1471-2164-10-33

Klasson, L, Walker, T, Sebaihia, M, Sanders, MJ, Quail, MA, Lord, A, Sanders, S, Earl, J, O’Neill, SL, Thomson, N, et al. 2008. Genome evolution of *Wolbachia* strain wPip from the *Culex pipiens* group. Molecular Biology and Evolution 25: 1877–1887.

Klasson, L, Westberg, J, Sapountzis, P, Naslund, K, Lutnaes, Y, Darby, AC, Veneti, Z, Chen, L, Braig, HR, Garrett, R, et al. 2009b. The mosaic genome structure of the *Wolbachia* wRi strain infecting *Drosophila* simulans. Proceedings of the National Academy of Sciences 106: 5725–5730. doi: 10.1073/pnas.0810753106

Kurtz, S, Phillippy, A, Delcher, AL, Smoot, M, Shumway, M, Antonescu, C, Salzberg, SL 2004. Versatile and open software for comparing large genomes. Genome Biology 5: R12. doi: 10.1186/gb-2004-5-2-r12

Lagesen, K, Hallin, P, Rødland, EA, Staerfeldt, H-H, Rognes, T, Ussery, DW 2007. RNAmmer: consistent and rapid annotation of ribosomal RNA genes. Nucleic Acids Research 35: 3100–3108. doi: 10.1093/nar/gkm160

Layton, EM, On, J, Perlmutter, JI, Bordenstein, SR, Shropshire, JD 2019. Paternal Grandmother Age Affects the Strength of *Wolbachia*-Induced Cytoplasmic Incompatibility in *Drosophila melanogaster*. MBio 10: e01879–01819. doi: 10.1128/mBio.01879-19

Lefoulon, E, Clark, T, Borveto, F, Perriat-Sanguinet, M, Moulia, C, Slatko, BE, Gavotte, L 2020. Pseudoscorpion *Wolbachia* symbionts: diversity and evidence for a new supergroup S. BMC Microbiology 20: 188. doi: 10.1186/s12866-020-01863-y

LePage, DP, Metcalf, JA, Bordenstein, SR, On, J, Perlmutter, JI, Shropshire, JD, Layton, EM, Funkhouser-Jones, LJ, Beckmann, JF, Bordenstein, SR 2017. Prophage WO genes recapitulate and enhance *Wolbachia*-induced cytoplasmic incompatibility. Nature 543: 243–247. doi: 10.1038/nature21391

Li, H, Durbin, R 2009. Fast and accurate short read alignment with Burrows-Wheeler transform. Bioinformatics 25: 1754–1760. doi: 10.1093/bioinformatics/btp324

Li, M, Brooks, CL, Wu-Baer, F, Chen, D, Baer, R, Gu, W 2003. Mono- versus polyubiquitination: differential control of p53 fate by Mdm2. Science 302: 1972–1975. doi: 10.1126/science.1091362

Li, W, Cowley, A, Uludag, M, Gur, T, McWilliam, H, Squizzato, S, Park, YM, Buso, N, Lopez, R 2015. The EMBL-EBI bioinformatics web and programmatic tools framework. Nucleic Acids Research 43: W580–584. doi: 10.1093/nar/gkv279

Lindsey, ARI, Rice, DW, Bordenstein, SR, Brooks, AW, Bordenstein, SR, Newton, ILG 2018. Evolutionary Genetics of Cytoplasmic Incompatibility Genes cifA and cifB in Prophage WO of *Wolbachia*. Genome Biology and Evolution 10: 434–451. doi: 10.1093/gbe/evy012

Lo, N, Paraskevopoulos, C, Bourtzis, K, O’Neill, SL, Werren, JH, Bordenstein, SR, Bandi, C 2007. Taxonomic status of the intracellular bacterium *Wolbachia pipientis*. International Journal of Systematic and Evolutionary 57: 654–657.

Lowe, TM, Eddy, SR 1997. tRNAscan-SE: a program for improved detection of transfer RNA genes in genomic sequence. Nucleic Acids Research 25: 955–964.

Madhav, M, Parry, R, Morgan, JAT, James, P, Asgari, S 2020. *Wolbachia* Endosymbiont of the Horn Fly (*Haematobia irritans irritans*): a Supergroup A Strain with Multiple Horizontally Acquired Cytoplasmic Incompatibility Genes. Applied and Environmental Microbiology 86: e02589–02519. doi: 10.1128/AEM.02589-19

Mardanov, AV, Ravin, NV 2006. Functional characterization of the repA replication gene of linear plasmid prophage N15. Research in Microbiology 157: 176–183. doi: https://doi.org/10.1016/j.resmic.2005.06.008

Martinez, J, Klasson, L, Welch, JJ, Jiggins, FM 2020. Life and death of selfish genes: comparative genomics reveals the dynamic evolution of cytoplasmic incompatibility. Molecular Biology and Evolution. doi: 10.1093/molbev/msaa209

Martinez, J, Longdon, B, Bauer, S, Chan, YS, Miller, WJ, Bourtzis, K, Teixeira, L, Jiggins, FM 2014. Symbionts commonly provide broad spectrum resistance to viruses in insects: a comparative analysis of *Wolbachia* strains. PLOS Pathogens 10: e1004369. doi: 10.1371/journal.ppat.1004369

Martinez, J, Ok, S, Smith, S, Snoeck, K, Day, JP, Jiggins, FM 2015. Should Symbionts Be Nice or Selfish? Antiviral Effects of *Wolbachia* Are Costly but Reproductive Parasitism Is Not. PLOS Pathogens 11: e1005021. doi: 10.1371/journal.ppat.1005021

McGraw, EA, Merritt, DJ, Droller, JN, O’Neill, SL 2001. *Wolbachia*-mediated sperm modification is dependent on the host genotype in *Drosophila*. Proceedings of the Royal Society B: Biological Sciences 268: 2565–2570. doi: 10.1098/rspb.2001.1839

McKenna, A, Hanna, M, Banks, E, Sivachenko, A, Cibulskis, K, Kernytsky, A, Garimella, K, Altshuler, D, Gabriel, S, Daly, M, et al. 2010. The Genome Analysis Toolkit: a MapReduce framework for analyzing next-generation DNA sequencing data. Genome Research 20: 1297–1303. doi: 10.1101/gr.107524.110

Merçot, H, Charlat, S 2004. *Wolbachia* infections in *Drosophila melanogaster* and *D. simulans*: polymorphism and levels of cytoplasmic incompatibility. Genetica 120: 51–59.

Mortier-Barriere, I, Velten, M, Dupaigne, P, Mirouze, N, Pietrement, O, McGovern, S, Fichant, G, Martin, B, Noirot, P, Le Cam, E, et al. 2007. A key presynaptic role in transformation for a widespread bacterial protein: DprA conveys incoming ssDNA to RecA. Cell 130: 824–836. doi: 10.1016/j.cell.2007.07.038

Newton, IL, Clark, ME, Kent, BN, Bordenstein, SR, Qu, J, Richards, S, Kelkar, YD, Werren, JH 2016. Comparative Genomics of Two Closely Related *Wolbachia* with Different Reproductive Effects on Hosts. Genome Biology and Evolution 8: 1526–1542. doi: 10.1093/gbe/evw096

Nikoh, N, Tsuchida, T, Maeda, T, Yamaguchi, K, Shigenobu, S, Koga, R, Fukatsu, T 2018. Genomic Insight into Symbiosis-Induced Insect Color Change by a Facultative Bacterial Endosymbiont, “*Candidatus* Rickettsiella viridis”. MBio 9: e00890–00818. doi: 10.1128/mBio.00890-18

Pati, A, Ivanova, NN, Mikhailova, N, Ovchinnikova, G, Hooper, SD, Lykidis, A, Kyrpides, NC 2010. GenePRIMP: a gene prediction improvement pipeline for prokaryotic genomes. Nature Methods 7: 455–457. doi: 10.1038/nmeth.1457

Peng, Y, Leung, HC, Yiu, SM, Chin, FY 2012. IDBA-UD: a de novo assembler for single-cell and metagenomic sequencing data with highly uneven depth. Bioinformatics 28: 1420–1428. doi: 10.1093/bioinformatics/bts174

Perlmutter, JI, Meyers, JE, Bordenstein, SR 2020. Transgenic Testing Does Not Support a Role for Additional Candidate Genes in *Wolbachia* Male Killing or Cytoplasmic Incompatibility. mSystems 5: e00658–00619. doi: 10.1128/mSystems.00658-19

Poinsot, D, Bourtzis, K, Markakis, G, Savakis, C, Mercot, H 1998. *Wolbachia* transfer from *Drosophila melanogaster* into *D. simulans*: Host effect and cytoplasmic incompatibility relationships. Genetics 150: 227–237.

Poinsot, D, Charlat, S, Mercot, H 2003. On the mechanism of *Wolbachia*-induced cytoplasmic incompatibility: confronting the models with the facts. Bioessays 25: 259–265. doi: 10.1002/bies.10234

Reveillaud, J, Bordenstein, SR, Cruaud, C, Shaiber, A, Esen, ÖC, Weill, M, Makoundou, P, Lolans, K, Watson, AR, Rakotoarivony, I, et al. 2019. The *Wolbachia* mobilome in *Culex pipiens* includes a putative plasmid. Nat Commun 10: 1051. doi: 10.1038/s41467-019-08973-w

Reynolds, KT, Hoffmann, AA 2002. Male age, host effects and the weak expression or non-expression of cytoplasmic incompatibility in *Drosophila* strains infected by maternally transmitted *Wolbachia*. Genetics Research 80: 79–87.

Rice, DW, Sheehan, KB, Newton, ILG 2017. Large-Scale Identification of *Wolbachia pipientis* Effectors. Genome Biology and Evolution 9: 1925–1937. doi: 10.1093/gbe/evx139

Rutherford, K, Parkhill, J, Crook, J, Horsnell, T, Rice, P, Rajandream, MA, Barrell, B 2000. Artemis: sequence visualization and annotation. Bioinformatics 16: 944–945.

Sasaki, T, Massaki, N, Kubo, T 2005. *Wolbachia* variant that induces two distinct reproductive phenotypes in different hosts. Heredity 95: 389–393.

Scholz, M, Albanese, D, Tuohy, K, Donati, C, Segata, N, Rota-Stabelli, O 2020. Large scale genome reconstructions illuminate *Wolbachia* evolution. Nat Commun 11: 5235. doi: 10.1038/s41467-020-19016-0

Schön, I, Kamiya, T, Van den Berghe, T, Van den Broecke, L, Martens, K 2019. Novel *Cardinium* strains in non-marine ostracod (Crustacea) hosts from natural populations. Molecular Phylogenetics and Evolution 130: 406–415. https://doi.org/10.1016/j.ympev.2018.09.008

Sheehan, KB, Martin, M, Lesser, CF, Isberg, RR, Newton, ILG 2016. Identification and Characterization of a Candidate *Wolbachia pipientis* Type IV Effector That Interacts with the Actin Cytoskeleton. MBio 7: e00622–00616. doi: 10.1128/mBio.00622-16

Shropshire, JD, Bordenstein, SR 2019. Two-By-One model of cytoplasmic incompatibility: Synthetic recapitulation by transgenic expression of cifA and cifB in *Drosophila*. PLoS Genetics 15: e1008221. doi: 10.1371/journal.pgen.1008221

Shropshire, JD, Kalra, M, Bordenstein, SR 2020a. Evolution-guided mutagenesis of the cytoplasmic incompatibility proteins: Identifying CifA’s complex functional repertoire and new essential regions in CifB. PLOS Pathogens 16: e1008794. doi: 10.1371/journal.ppat.1008794

Shropshire, JD, Leigh, B, Bordenstein, SR 2020b. Symbiont-mediated cytoplasmic incompatibility: what have we learned in 50 years? Elife 9: e61989. doi: 10.7554/eLife.61989

Shropshire, JD, Leigh, B, Bordenstein, SR, Duplouy, A, Riegler, M, Brownlie, JC, Bordenstein, SR 2019. Models and Nomenclature for Cytoplasmic Incompatibility: Caution over Premature Conclusions – A Response to Beckmann et al. Trends in Genetics 35: 397–399. https://doi.org/10.1016/j.tig.2019.03.004

Shropshire, JD, On, J, Layton, EM, Zhou, H, Bordenstein, SR 2018. One prophage WO gene rescues cytoplasmic incompatibility in *Drosophila melanogaster*. Proceedings of the National Academy of Sciences 115: 4987. doi: 10.1073/pnas.1800650115

Simpson, JT, Wong, K, Jackman, SD, Schein, JE, Jones, SJ, Birol, I 2009. ABySS: a parallel assembler for short read sequence data. Genome Research 19: 1117–1123. doi: 10.1101/gr.089532.108

Sinha, A, Li, Z, Sun, L, Carlow, CKS 2019. Complete genome sequence of the *Wolbachia* wAlbB endosymbiont of *Aedes albopictus*. Genome Biology and Evolution 11: 706–720. doi: 10.1093/gbe/evz025

Smeets, LC, Bijlsma, JJ, Kuipers, EJ, Vandenbroucke-Grauls, CM, Kusters, JG 2000. The dprA gene is required for natural transformation of Helicobacter pylori. FEMS Immunol Med Microbiol 27: 99–102.

Stamatakis, A 2006. RAxML-VI-HPC: maximum likelihood-based phylogenetic analyses with thousands of taxa and mixed models. Bioinformatics 22: 2688–2690. doi: 10.1093/bioinformatics/btl446

Stewart, AC, Osborne, B, Read, TD 2009. DIYA: a bacterial annotation pipeline for any genomics lab. Bioinformatics 25: 962–963. doi: 10.1093/bioinformatics/btp097

Stouthamer, CM, Kelly, SE, Mann, E, Schmitz-Esser, S, Hunter, MS 2019. Development of a multi-locus sequence typing system helps reveal the evolution of *Cardinium hertigii*, a reproductive manipulator symbiont of insects. BMC Microbiology 19: 266. doi: 10.1186/s12866-019-1638-9

Sutton, ER, Harris, SR, Parkhill, J, Sinkins, SP 2014. Comparative genome analysis of *Wolbachia* strain wAu. BMC Genomics 15: 928. doi: 10.1186/1471-2164-15-928

Takata, T, Ando, T, Israel, DA, Wassenaar, TM, Blaser, MJ 2005. Role of dprA in transformation of *Campylobacter jejuni*. FEMS Microbiology Letters 252: 161–168. doi: 10.1016/j.femsle.2005.08.052

Teixeira, L, Ferreira, A, Ashburner, M 2008. The bacterial symbiont *Wolbachia* induces resistance to RNA viral infections in *Drosophila melanogaster*. PLoS Biology 6: e2.

Teixeira, LK, Reed, SI 2013. Ubiquitin ligases and cell cycle control. Annu Rev Biochem 82: 387–414. doi: 10.1146/annurev-biochem-060410-105307

Toomey, ME, Frydman, HM 2014. Extreme divergence of *Wolbachia* tropism for the stem-cell-niche in the *Drosophila* testis. PLOS Pathogens 10: e1004577. doi: 10.1371/journal.ppat.1004577

Toomey, ME, Panaram, K, Fast, EM, Beatty, C, Frydman, HM 2013. Evolutionarily conserved *Wolbachia*-encoded factors control pattern of stem-cell niche tropism in *Drosophila* ovaries and favor infection. Proceedings of the National Academy of Sciences of the United States of America 110: 10788–10793. doi: 10.1073/pnas.1301524110

Tram, U, Sullivan, W 2002. Role of delayed nuclear envelope breakdown and mitosis in *Wolbachia*-induced cytoplasmic incompatibility. Science 296: 1124–1126.

Tsuchida, T, Koga, R, Horikawa, M, Tsunoda, T, Maoka, T, Matsumoto, S, Simon, J-C, Fukatsu, T 2010. Symbiotic Bacterium Modifies Aphid Body Color. Science 330: 1102. doi: 10.1126/science.1195463

Uehara, R, Nozawa, R-s, Tomioka, A, Petry, S, Vale, RD, Obuse, C, Goshima, G 2009. The augmin complex plays a critical role in spindle microtubule generation for mitotic progression and cytokinesis in human cells. Proceedings of the National Academy of Sciences of the United States of America 106: 6998–7003. doi: 10.1073/pnas.0901587106

Veneti, Z, Clark, ME, Karr, TL, Savakis, C, Bourtzis, K 2004. Heads or tails: host-parasite interactions in the *Drosophila*-*Wolbachia* system. Applied Environmental Microbiology 70: 5366–5372. doi: 10.1128/AEM.70.9.5366-5372.2004 70/9/5366 [pii]

Veneti, Z, Clark, ME, Zabalou, S, Karr, TL, Savakis, C, Bourtzis, K 2003. Cytoplasmic incompatibility and sperm cyst infection in different *Drosophila*-*Wolbachia* associations. Genetics 164: 545–552.

Wang, GH, Sun, BF, Xiong, TL, Wang, YK, Murfin, KE, Xiao, JH, Huang, DW 2016. Bacteriophage WO Can Mediate Horizontal Gene Transfer in Endosymbiotic *Wolbachia* Genomes. Front Microbiol 7: 1867. doi: 10.3389/fmicb.2016.01867

Weldon, SR, Strand, MR, Oliver, KM 2013. Phage loss and the breakdown of a defensive symbiosis in aphids. Proceedings of the Royal Society B: Biological Sciences 280: 20122103. doi: 10.1098/rspb.2012.2103

Wernegreen, JJ, Moran, NA 2001. Vertical transmission of biosynthetic plasmids in aphid endosymbionts (*Buchnera*). Journal of Bacteriology 183: 785–790. doi: 10.1128/JB.183.2.785-790.2001

Werren, JH 1997. Biology of *Wolbachia*. Annual Review of Entomology 42: 587–609. doi: 10.1146/annurev.ento.42.1.587

Werren, JH, Baldo, L, Clark, ME 2008. *Wolbachia*: master manipulators of invertebrate biology. Nature Reviews Microbiology 6: 741–751. doi: 10.1038/nrmicro1969

Woolfit, M, Iturbe-Ormaetxe, I, McGraw, EA, O’Neill, SL 2009. An ancient horizontal gene transfer between mosquito and the endosymbiotic bacterium *Wolbachia pipientis*. Molecular Biology and Evolution 26: 367–374. doi: msn253 [pii] 10.1093/molbev/msn253

Wu, M, Sun, LV, Vamathevan, J, Riegler, M, Deboy, R, Brownlie, JC, McGraw, EA, Martin, W, Esser, C, Ahmadinejad, N, et al. 2004. Phylogenomics of the reproductive parasite *Wolbachia pipientis* wMel: a streamlined genome overrun by mobile genetic elements. PLoS Biology 2: E69. doi: 10.1371/journal.pbio.0020069

Yang, Z 2007. PAML 4: phylogenetic analysis by maximum likelihood. Molecular Biology and Evolution 24: 1586–1591. doi: 10.1093/molbev/msm088

Zabalou, S, Apostolaki, A, Pattas, S, Veneti, Z, Paraskevopoulos, C, Livadaras, I, Markakis, G, Brissac, T, Mercot, H, Bourtzis, K 2008. Multiple rescue factors within a Wolbachia strain. Genetics 178: 2145–2160. doi: 178/4/2145 [pii] 10.1534/genetics.107.086488

Zabalou, S, Charlat, S, Nirgianaki, A, Lachaise, D, Mercot, H, Bourtzis, K 2004. Natural *Wolbachia* infections in the *Drosophila yakuba* species complex do not induce cytoplasmic incompatibility but fully rescue the wRi modification. Genetics 167: 827–834. doi: 10.1534/genetics.103.015990 167/2/827 [pii]

Zchori-Fein, E, Perlman, SJ 2004. Distribution of the bacterial symbiont *Cardinium* in arthropods. Molecular Ecology 13: 2009–2016. doi: 10.1111/j.1365-294X.2004.02203.x MEC2203 [pii]

Zerbino, DR, Birney, E 2008. Velvet: algorithms for de novo short read assembly using de Bruijn graphs. Genome Research 18: 821–829. doi: 10.1101/gr.074492.107

Zheng, X, Zhang, D, Li, Y, Yang, C, Wu, Y, Liang, X, Liang, Y, Pan, X, Hu, L, Sun, Q, et al. 2019. Incompatible and sterile insect techniques combined eliminate mosquitoes. Nature 572: 56–61. doi: 10.1038/s41586-019-1407-9

Zug, R, Hammerstein, P 2012. Still a host of hosts for *Wolbachia*: Analysis of recent data suggests that 40% of terrestrial arthropod species are infected. PLoS ONE 7: e38544. doi: 10.1371/journal.pone.0038544

