## Supplementary figures for "Comparative genomics reveals factors associated with phenotypic expression of *Wolbachia*"

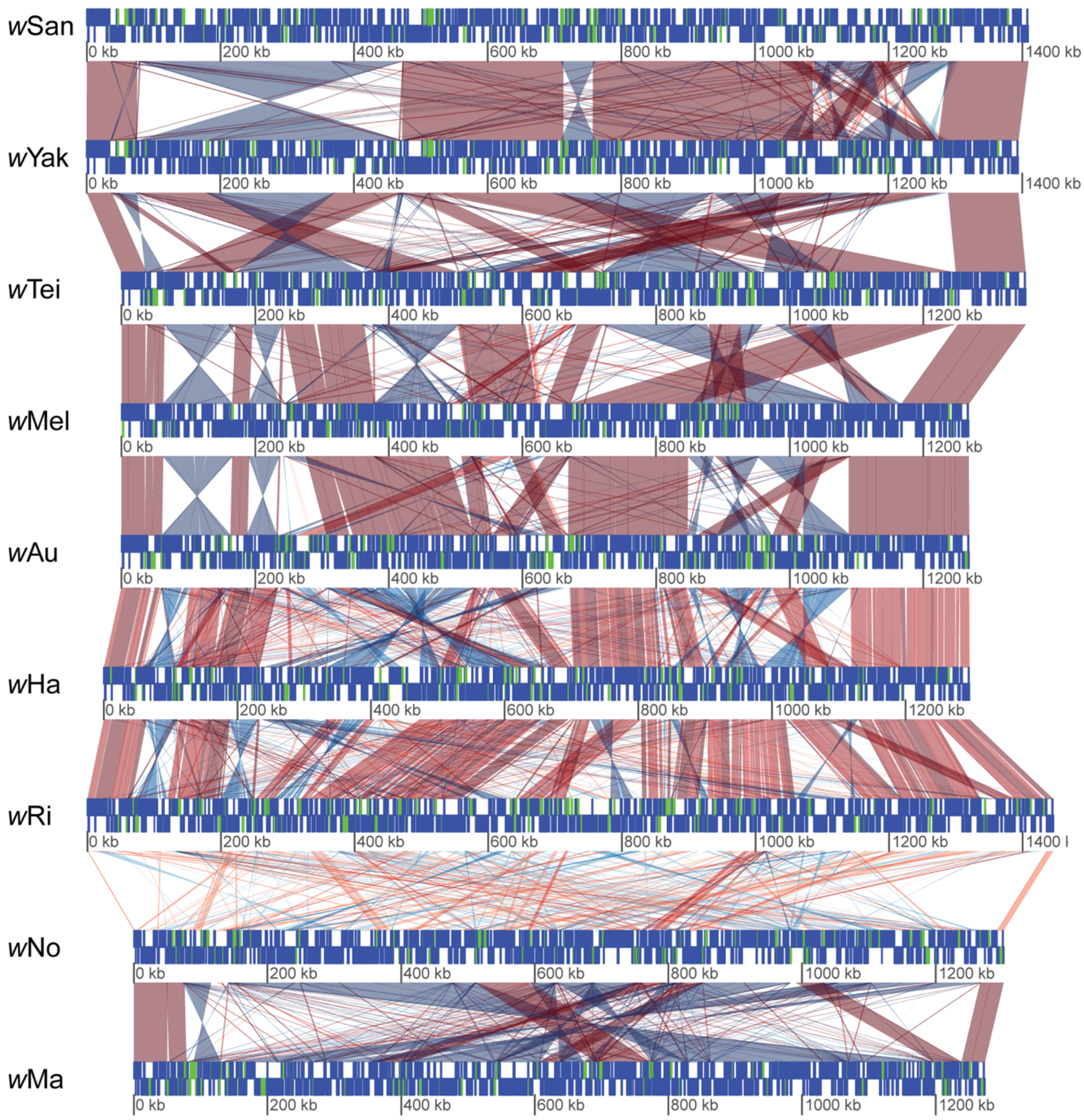


**Figure S1. Whole-genome alignment of the nine *Wolbachia* genomes included in the study.** Blue blocks symbolize functional genes and green blocks indicate pseudogenes. Similarity between sequences is indicated by the intensity of the red (forward macth) or blue (reverse match) lines, where darker is more similar.

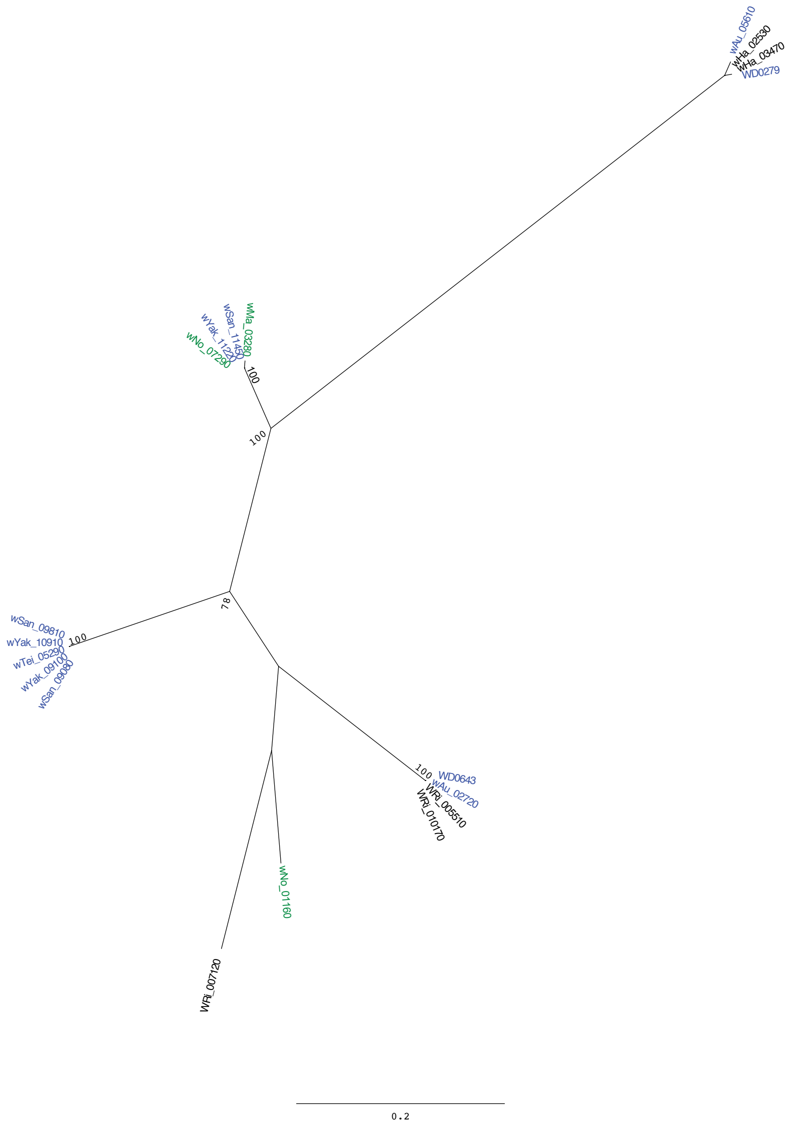


**Figure S2. Phylogenetic tree of a WO phage gene and its homologs in our genomes.** Maximum likelihood tree inferred with RAxML using 100 bootstraps. Only boostrap values above 70 are shown. Branch labels in blue indicate genes from the SYTMA genomes and branch labels in green indicate genes from the NoMa genomes. The branch with only blue lables contain the divergent *w*SYT phage copy.

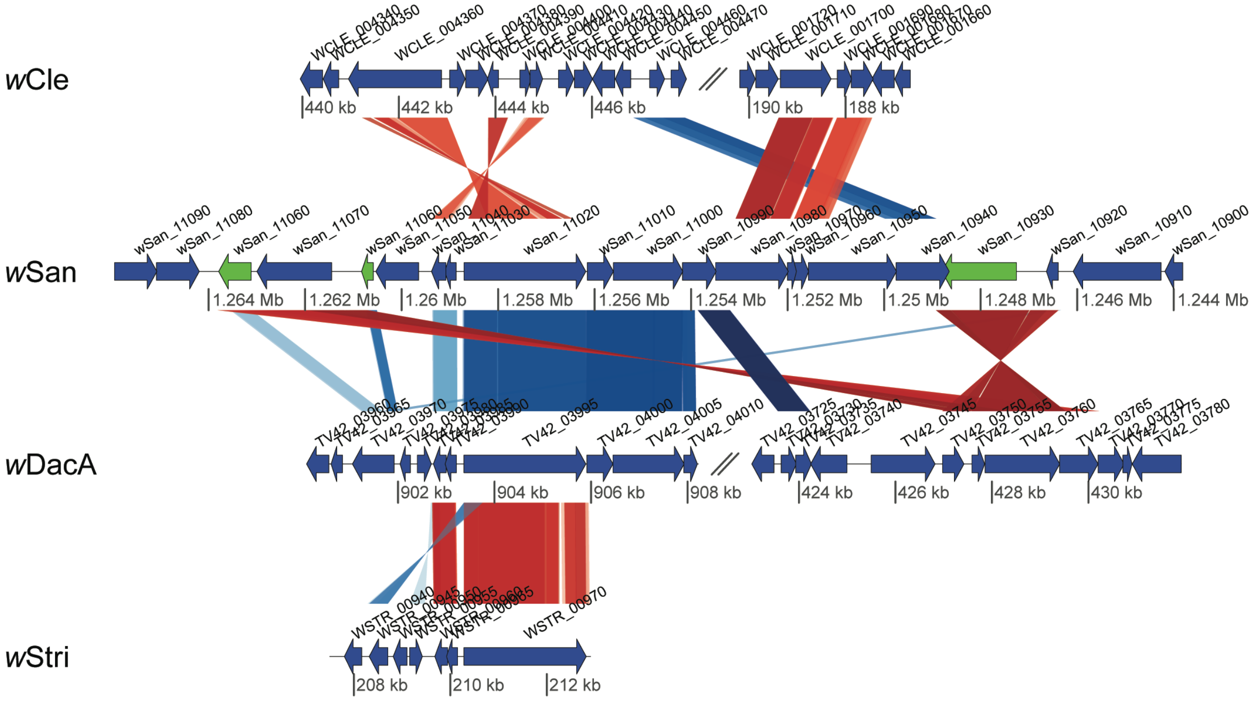


**Figure S3. Similarity between a twelve gene genomic region in *w*SYT and distantly related *Wolbachia* genomes.** Blue arrows symbolize functional genes and green arrows indicate pseudogenes. Similarity between sequences is indicated by the intensity of the red (forward macth) or blue (reverse match) lines, where darker is more similar

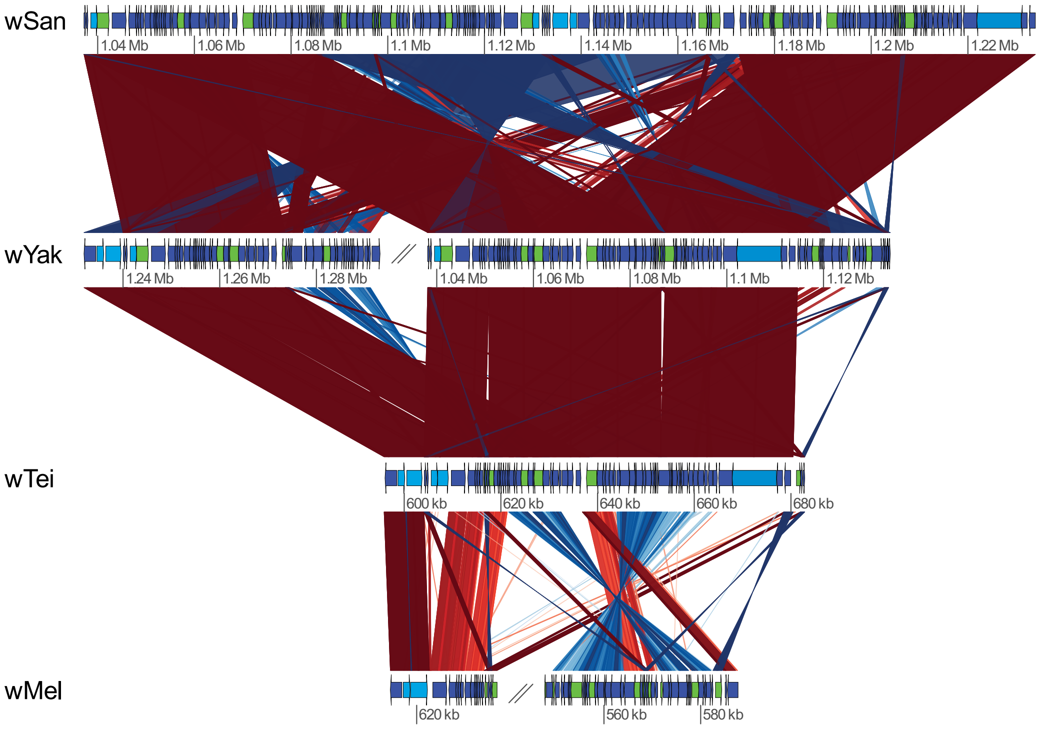


**Figure S4. Comparison of WO phage regions between SYTM genomes.** Blue arrows symbolize functional genes and green arrows indicate pseudogenes. Putative functional *cif*-homologs and the WD0513-homolog are marked in lighter blue. Similarity between sequences is indicated by the intensity of the red (forward match) or blue (reverse match) lines, where darker is more similar.

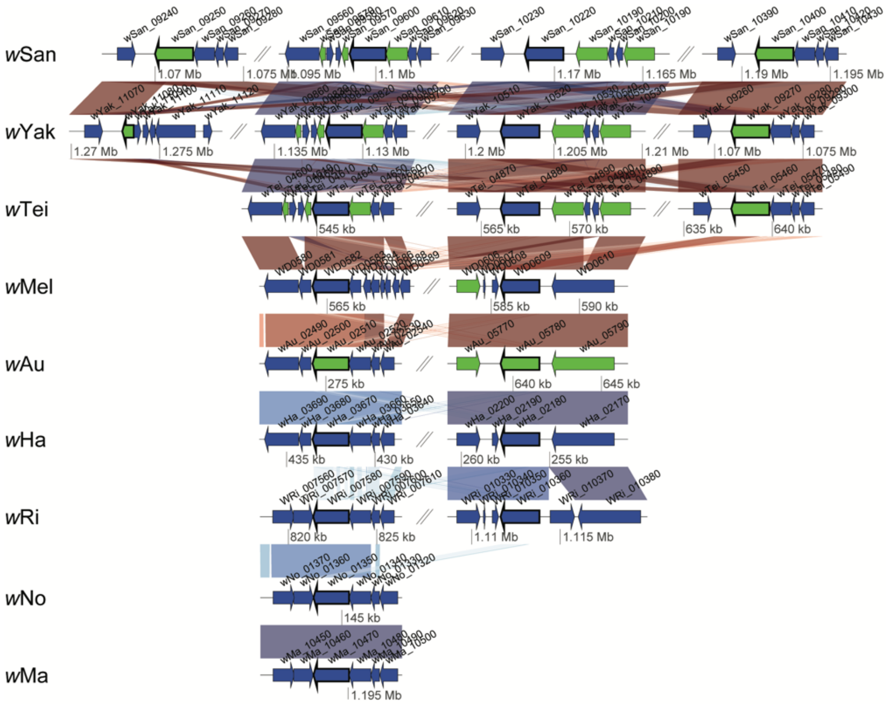


**Figure S5. Comparison of the phage *repA* gene and its flanking region**. Blue arrows symbolize functional genes and green arrows indicate pseudogenes. The *repA* genes are marked with a black border. Similarity between sequences is indicated by the intensity of the red (forward macth) or blue (reverse match) lines, where darker is more similar.

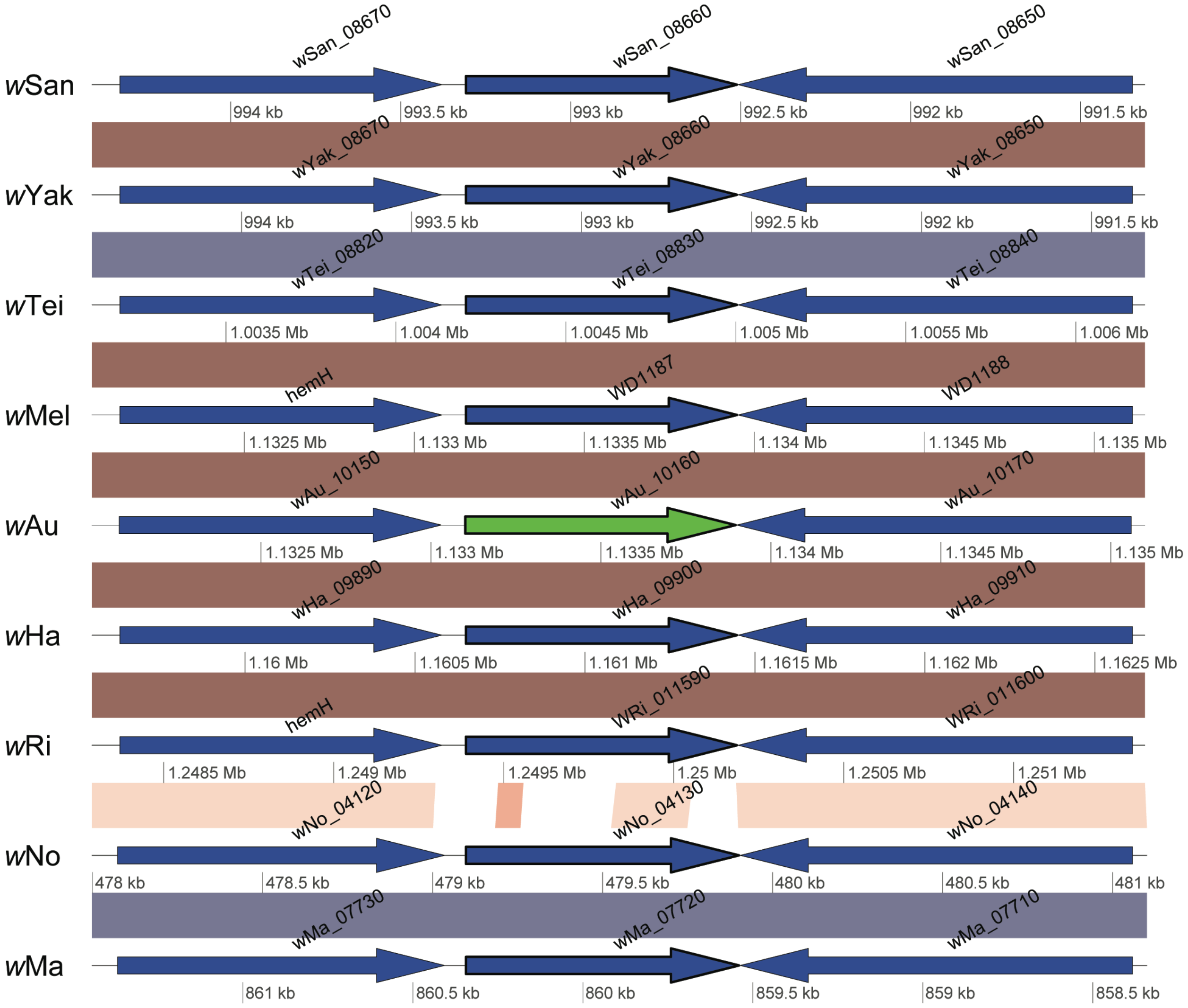


**Figure S6. Comparison of the genomic region containing the *w*Mel gene WD1187**. Blue arrows symbolize functional genes and green arrows indicate pseudogenes. WD1187-homologs are marked with a black border. Similarity between sequences is indicated by the intensity of the red (forward macth) or blue (reverse match) lines, where darker is more similar

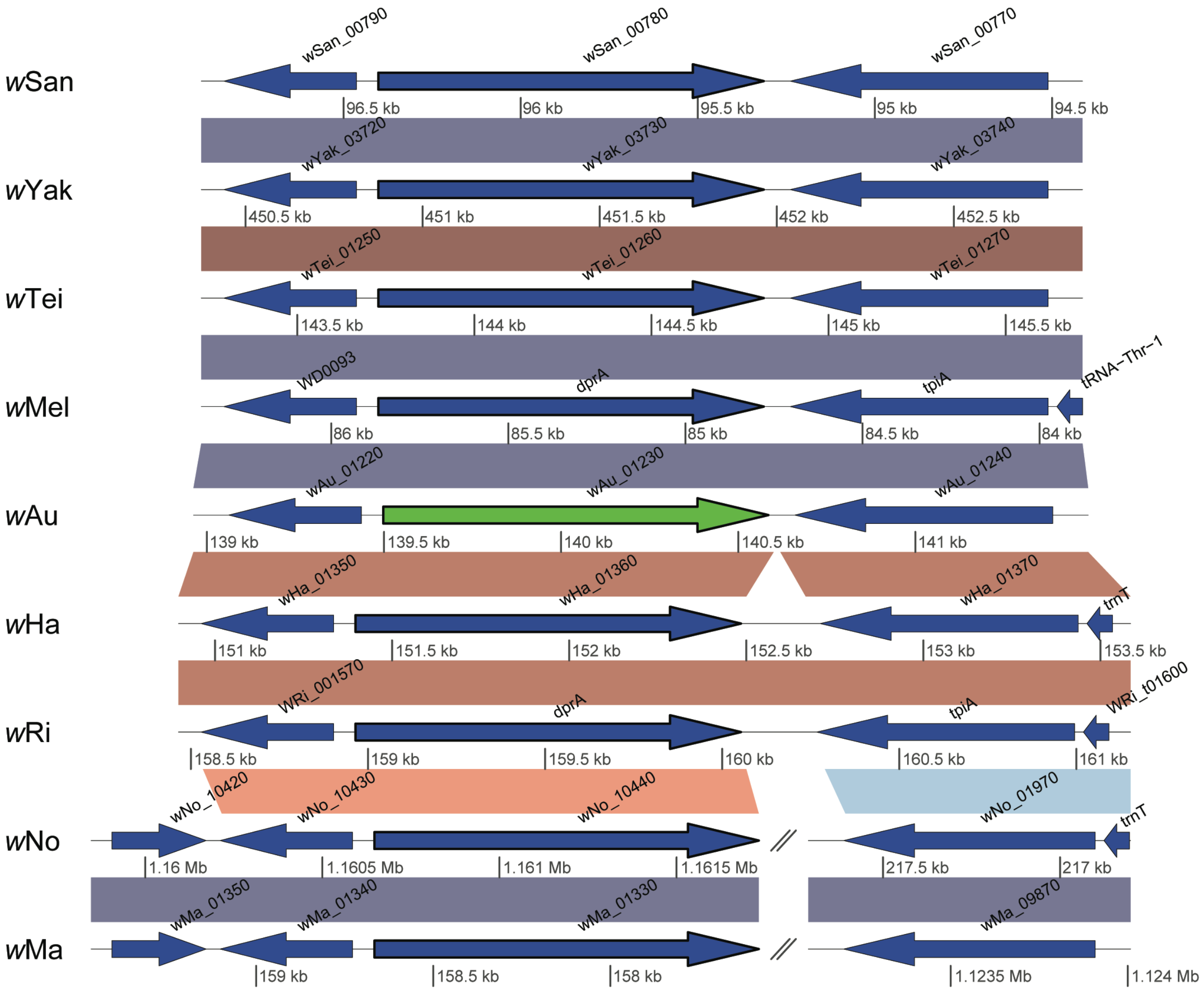


**Figure S7. Comparison of the genomic region containing *dprA* gene**. Blue arrows symbolize functional genes and green arrows indicate pseudogenes. *dprA*-homologs are marked with a black border. Similarity between sequences is indicated by the intensity of the red (forward macth) or blue (reverse match) lines, where darker is more similar
